# Effect of Ca^2+^ on the promiscuous target-protein binding mechanism of calmodulin

**DOI:** 10.1101/277327

**Authors:** Annie M. Westerlund, Lucie Delemotte

**Affiliations:** Science for Life Laboratory, Department of Applied Physics, KTH Royal Institute of Technology, Stockholm, Sweden

## Abstract

Calmodulin (CaM) is a calcium sensing protein that regulates the function of a large number of proteins, thus playing a crucial part in many cell signaling pathways. CaM has the ability to bind more than 300 different target peptides in a Ca^2+^-dependent manner, mainly through the exposure of hydrophobic residues. How CaM can bind a large number of targets while retaining some selectivity is a fascinating open question.

Here, we explore the mechanism of CaM selective promiscuity for selected target proteins. Analyzing enhanced sampling molecular dynamics simulations of Ca^2+^-bound and Ca^2+^-free CaM via spectral clustering has allowed us to identify distinct conformational states, characterized by interhelical angles, secondary structure determinants and the solvent exposure of specific residues. We searched for indicators of conformational selection by mapping solvent exposure of residues in these conformational states to contacts in structures of CaM/target peptide complexes. We thereby identified CaM states involved in various binding classes arranged along a depth binding gradient. Binding Ca^2+^ modifies the accessible hydrophobic surface of the two lobes and allows for deeper binding. Apo CaM indeed shows shallow binding involving predominantly polar and charged residues. Furthermore, binding to the C-terminal lobe of CaM appears selective and involves specific conformational states that can facilitate deep binding to target proteins, while binding to the N-terminal lobe appears to happen through a more flexible mechanism. Thus the long-ranged electrostatic interactions of the charged residues of the N-terminal lobe of CaM may initiate binding, while the short-ranged interactions of hydrophobic residues in the C-terminal lobe of CaM may account for selectivity.

This work furthers our understanding of the mechanism of CaM binding and selectivity to different target proteins and paves the way towards a comprehensive model of CaM selectivity.

**Author summary:** Calmodulin is a protein involved in the regulation of a variety of cell signaling pathways. It acts by making usually calcium-insensitive proteins sensitive to changes in the calcium concentration inside the cell. Its two lobes bind calcium and allow the energetically unfavorable exposure of hydrophobic residues to the aqueous environment which can then bind target proteins. The mechanisms behind the simultaneous specificity and variation of target protein binding is yet unknown but will aid understanding of the calcium-signaling and regulation that occur in many of our cellular processes.

Here, we used molecular dynamics simulations and data analysis techniques to investigate what effect calcium has on the binding modes of calmodulin. The simulations and analyses allow us to observe and differentiate specific states. One domain of calmodulin is shown to be selective with binding involving short-distance interactions between hydrophobic residues, while the other binds target proteins through a more flexible mechanism involving long-distance electrostatic interactions.

## Introduction

Calmodulin (CaM), Fig 1.a-b, is a promiscuous Ca^2+^-sensing protein that plays part in many physiologically important cellular processes [1]. Its two lobes, connected by a flexible linker, have one beta sheet and two EF-hand motifs each, Fig 1.a-c. The EF-hand binds a Ca^2+^ ion which induces tertiary structure rearrangements of lobe helices, exposing hydrophobic residues [2]. This allows CaM to bind and regulate a myriad of target proteins such as ion channels, kinases and G-protein coupled receptors. The Ca^2+^-signaling and olfactory transduction pathways (Fig 1.d-e) are two examples of cell-signaling pathways where CaM is involved. In the Ca^2+^-signaling pathway, CaM activates and regulates the myosin light chain kinase IV (MLCK) and calcineurin (CaN), among others [3]. Through this, CaM plays part in regulating a variety of biological processes including metabolism, proliferation and learning. In olfactory transduction, CaM’s role is instead to inhibit the CNG channel and activate the Ca^2+^/CaM-dependent protein kinase (CAMK) which inhibits the adenylate cyclase 3 (ADCY3).

**Fig 1.**
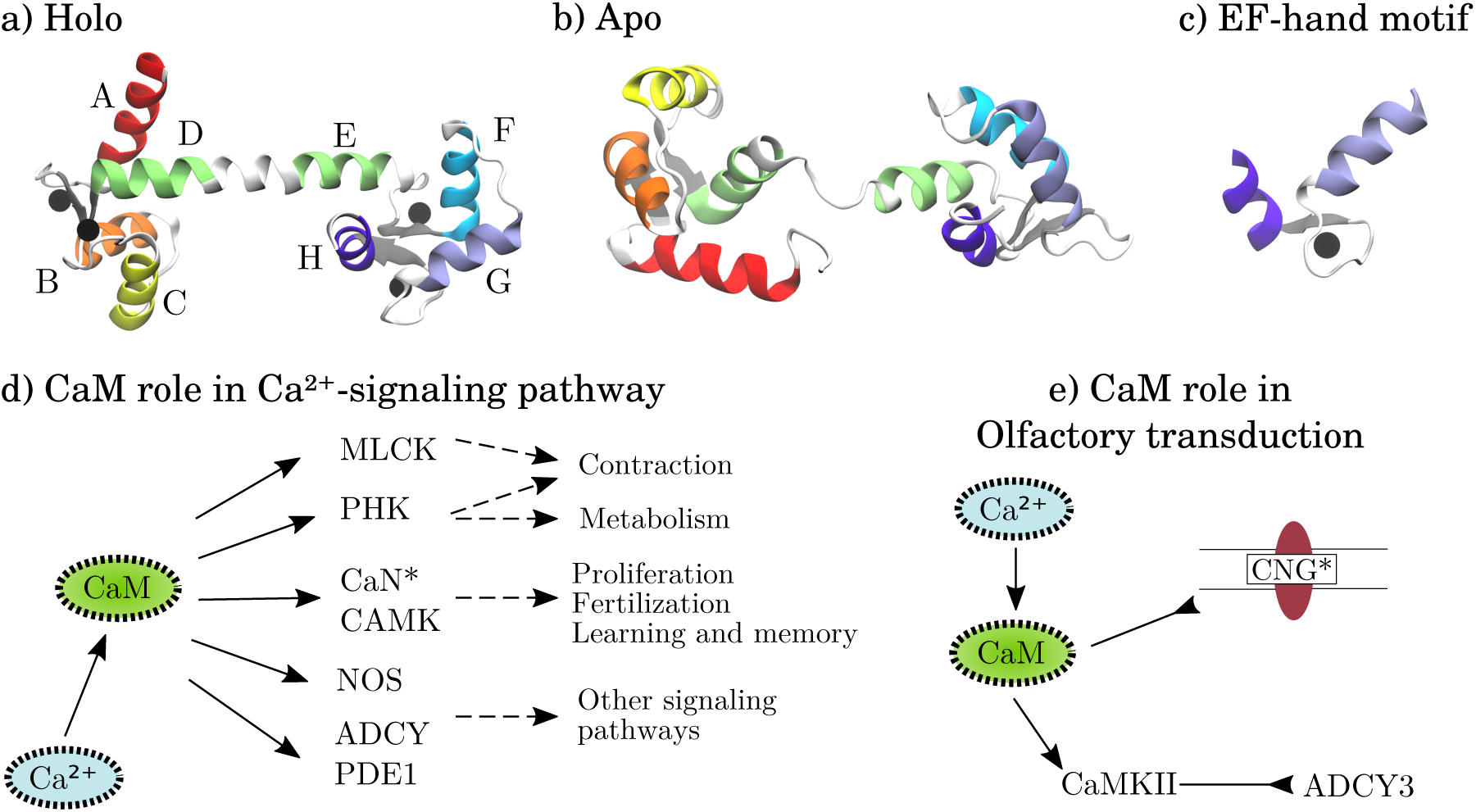
The molecular structure of calmodulin and pathways where calmodulin acts as protein regulator. Molecular structures of a) holo and b) apo calmodulin. The helices are marked according to their canonical labeling. Ca^2+^ ions are represented as black spheres and the beta sheets are marked with gray color. c) The EF-hand motif. d) The role of CaM in the Ca^2+^-signaling pathway. CaM activates the myosin light chain kinase IV (MLCK) and phosphorylase kinase (PHK), calcineurin (CaN), Ca^2+^/calmodulin-dependent protein kinase (CAMK), nitric oxide synthase 1 (NOS), adenylate cyclase 1 (ADCY) and phosphodiesterase 1A (PDE1). This affects downstream processes such as contraction, metabolism, proliferation, learning etc. e) The role of CaM in olfactory transduction. CaM inhibits the cyclic nucleotide-gated (CNG) channel and activates Ca^2+^/calmodulin-dependent protein kinase II (CaMKII). CaMKII then inhibits adenylate cyclase 3 (ADCY3). Proteins marked by a star are included in our CaM binding study. The pathways in d) and e) are adapted from KEGG [3].

CaM’s promiscuity can be mainly ascribed to its ubiquity and interactions to specific target proteins depend partly on local cell environments such as availability of target proteins. Moreover, high affinity binding to target proteins is partly due to the flexible linker that allows wrapping around target proteins but also stems from the structural properties of the two lobes, thus linked to the pockets formed by their hydrophobic interfaces [4]. Here, we study the interactions between CaM and various target proteins, including proteins that are involved in the Ca^2+^-signaling and olfactory transduction pathways.

Ca^2+^-independent CaM regulation, with Ca^2+^-free CaM (apo CaM) regulat-ing the target protein may occur, although not as frequently as Ca^2+^-dependent CaM regulation. The IQ binding motif, named after the first two conserved residues of the motif, is often associated to apo-CaM binding while 1-5-10 or 1-5-8-14 motifs, named after the position of conserved hydrophobic residues, are associated with Ca^2+^-bound CaM (holo CaM) binding. [5–7] These motifs are, however, lacking the complexity to completely distinguish holo-CaM and apo-CaM binding, as well as different types of binding. An example is seen in Fig 2, where apo and holo CaM C-terminal domain (C-CaM) expose different hydrophobic interfaces, but both bind the voltage-gated sodium channel Na_V_1.2 at the same IQ binding motif. Holo CaM often interacts with a higher affinity, but in some cases, such as the IQ motif of Na_v_1.2, apo-CaM binding is more favorable than holo-CaM binding [8, 9].

**Fig 2.**
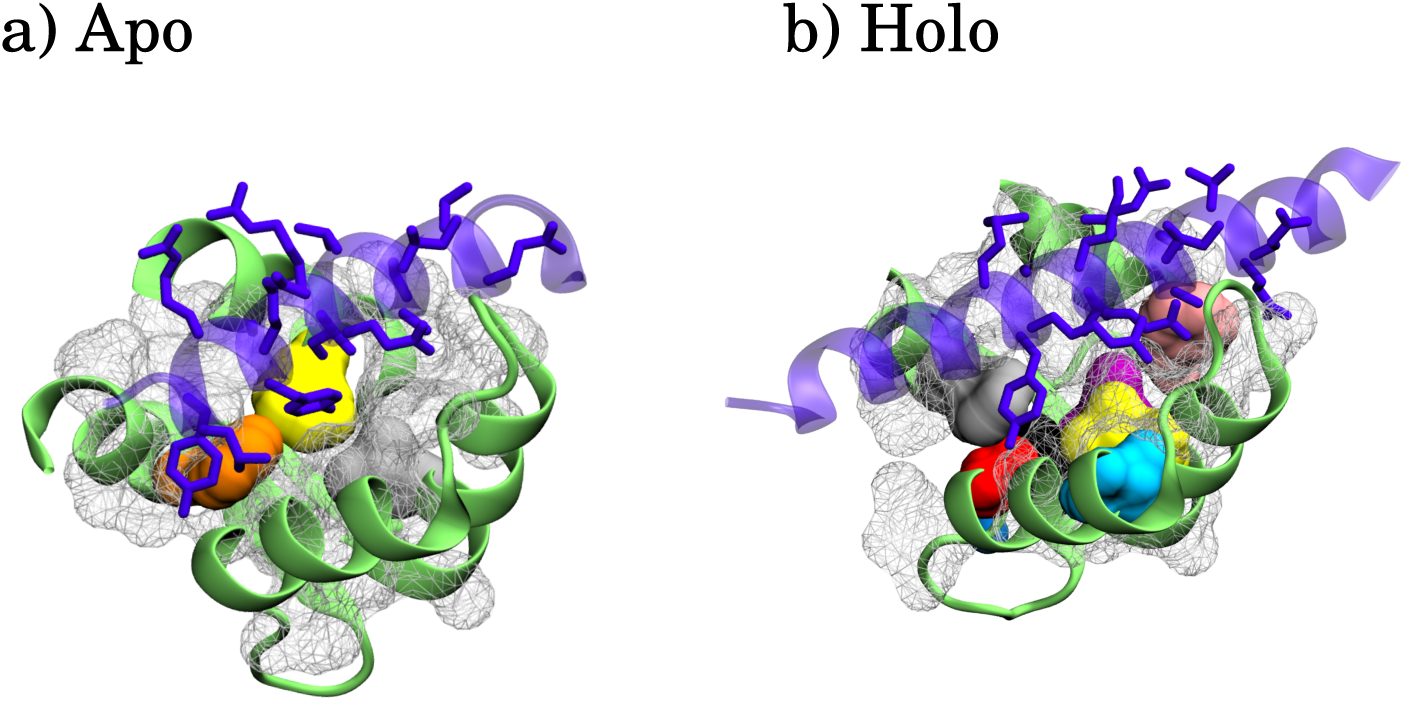
Calmodulin C-term domain binding the voltage-gated sodium channel Na_V_1.2. a) Ca^2+^-free state (apo) and b) Ca^2+^-bound state (holo). The structures have PDB accession numbers a) 2KXW and b) 4JPZ. The target protein is violet with side chains depicted as licorice. Hydrophobic residues are visualized in a white mesh. Key hydrophobic residues found in this paper’s analysis are highlighted in different colors. The residues are shown in S3 Fig and S1 Fig.

Both CaM dynamics and motif-dependent target-protein binding have been extensively studied via both experiments [2, 8, 10–25] and simulations. Indeed, all-atom molecular dynamics (MD) simulations allow for a detailed description with molecular insights. Early MD simulations of the holo CaM conformational ensemble have shown the holo-CaM linker to be flexible [26–28] and indicated structural destabilization when removing Ca^2+^ from one site in the N-term domain [29]. Moreover, CaM target-protein binding was approached by studying the dynamics as well as the thermodynamics of CaM or CaM complexes [4, 26, 30], with insights into binding of specific target proteins using both regular MD and metadynamics [30]. However, little is known about the conformations of the binding interfaces and different binding modes of the two lobes, as well as their relation to protein-unbound CaM conformational ensemble. Part of this was addressed through the analysis of multiple aggregated MD simulations of C-CaM using Markov state modeling (MSM), in which target protein binding was proposed to occur by C-CaM adopting the bound conformation before binding to the protein [31]. However, the analysis did not cover the differences of CaM N-terminal domain (N-CaM) and C-CaM dynamics, mechanisms and binding modes, nor did the study explore the possibility of different binding mechanisms linked to different binding modes.

Binding a ligand can be conceptualized using two frameworks: conformational selection and induced fit. In pure conformational selection, the apo protein adopts a holo-like state before binding [32]. In pure induced fit, the ligand binds in a typical apo state that is not ideal, which causes subsequent rearrangements before reaching a high-affinity holo state [33]. In reality, binding likely involves a combination of the two mechanisms. However, spectroscopy experiments, as well as extensive MD simulations, may shed light on which mechanism is dominating. If the apo protein samples the holo-like state, conformational selection is typically assumed to be dominating, otherwise induced fit is assumed [34].

Here, we analyze thermally enhanced MD simulations of calmodulin with different Ca^2+^-occupancy and use spectral clustering to elucidate calmodulin selective promiscuity. We search for indicators of conformational selection by mapping solvent exposure of residues from sampled states (clusters) to contacts of already existing structures of CaM/peptide complexes. Moreover, we gain knowledge about the characteristics behind different binding modes of the two lobes, as well as the difference between holo and apo binding modes.

## Materials and Methods

### Calmodulin system preparation and equilibration

For this project, we considered four different binding states: holo and apo calmodulin, as well as Ca^2+^ bound only in the N-term (N-holo) and Ca^2+^ bound only in the C-term (C-holo), Table 1. The simulations of N-holo, C-holo and holo CaM used structure 3CLN [35], while the apo simulations used structure 1LKJ [36]. N-holo was generated by removing Ca^2+^ from C-CaM and C-holo by removing Ca^2+^ from N-CaM from the holo structure. The systems were built using Charmm-gui [37, 38], where the protein was solvated in a box of about 21000 water molecules. The systems were then ionized with 0.15 M NaCl. Charmm36 was chosen as force field [39], and TIP3P [40] as water model. The modified parameters of Charmm27 force field from Marinelli and Faraldo-Gomez [41] were used to parameterize Ca^2+^ ions.

**Table 1.** Systems and MD simulation details

| Ca<sup>2+</sup>-binding state | PDB ID | Number of atoms | Length [MD, T-REMD, REST] |
| --- | --- | --- | --- |
| **Holo** | 3CLN | 65620 | [3600, 570, 460] ns |
| **Apo** | 1LKJ | 66277 | [5800, 570, 420] ns |
| **Ca<sup>2+</sup> in N-term (N-holo)** | 3CLN | 65614 | [2400, 570, 420] ns |
| **Ca<sup>2+</sup> in C-term (C-holo)** | 3CLN | 65614 | [2100, 570, 350] ns |

5000 steps of minimization were carried out, followed by a 50 ps NVT ensemble equilibration with strong harmonic restraints on the protein atoms. The box was then scaled, relaxing pressure with Berendsen barostat [42], while gradually releasing the position restraints for 350 ps.

### Molecular dynamics simulation parameters

The MD parameters used in these simulations are extensively described elsewhere [43].

Calmodulin was simulated in an NPT ensemble with a 1 atm pressure and 2 fs time step. The short-ranged electrostatic interactions were modeled with a 1.2 nm cutoff where the switching function started at 1.0 nm, and the long-ranged ones with PME [44]. We used a Nose-Hoover thermostat [45], an isotropic Parinello-Rahman barostat [46], and constrained hydrogen bonds with LINCS [47].

The MD simulations were run at a constant temperature of 303.5 K. In addition to regular MD simulations, temperature enhanced simulations were performed; temperature replica exchange [48] (T-REMD) and replica exchange solute tempering [49–51] (REST). In T-REMD, parallel simulations of independent replicas are run at different temperatures. A random walk in temperature space is achieved by employing the Metropolis criterion periodically to accept coordinate exchanges between neighboring replicas. Conformations obtained at higher temperatures are propagated down to the lower temperatures through exchanges. Because barriers are more easily passed at higher temperatures, the efficiency is increased when the free energy landscape is rugged. However, the number of replicas needed to span a certain temperature interval scales with system size [52]. For this reason, T-REMD may not always be efficient. To alleviate this, REST only modifies the hamiltonian of the system for the solute (protein), and not the solvent [51]. However, the relative efficiencies of regular MD, T-REMD and REST depend on the ruggedness of the free energy landscape [43].

We used 25 replicas in both T-REMD and REST. The replicas of T-REMD spanned a temperature range of 299.13-326.09 K, while the REST replicas were simulated at temperatures between 300.0-545.0 K. The temperature ranges for REST and T-REMD were chosen using the “Temperature generator for REMD simulations” [53], considering only the protein for REST. Exchanges between neighboring replicas were attempted every 2 ps, where half of the replicas were involved in each attempt.

The REST simulations were performed with the Plumed 2.3b [54] plug-in patched with Gromacs version 5.1.2 [55], where the charge of the atoms in the hot region were scaled, as well as the interactions between the two regions and the proper dihedral angles [56]. Analysis was performed using the replica at the lowest temperature. and was carried out on the protein heavy atoms. The first four residues in the apo structure were removed, because those are missing in the 3CLN structure.

The apo CaM ensemble is more diffusive and generally well sampled by regular MD, while holo CaM free energy landscape is more hierarchical, and thus sampled more efficiently by temperature enhanced methods [43]. For this reason, more MD data is used in the apo analysis while more REST data is used for holo, Table 1. The three methods may sample different regions with varying efficiency, which is why data from all three simulations is used.

### Identifying states through spectral clustering

Fig 3 illustrates the steps carried out to analyze the MD simulation trajectories.

**Fig 3.**
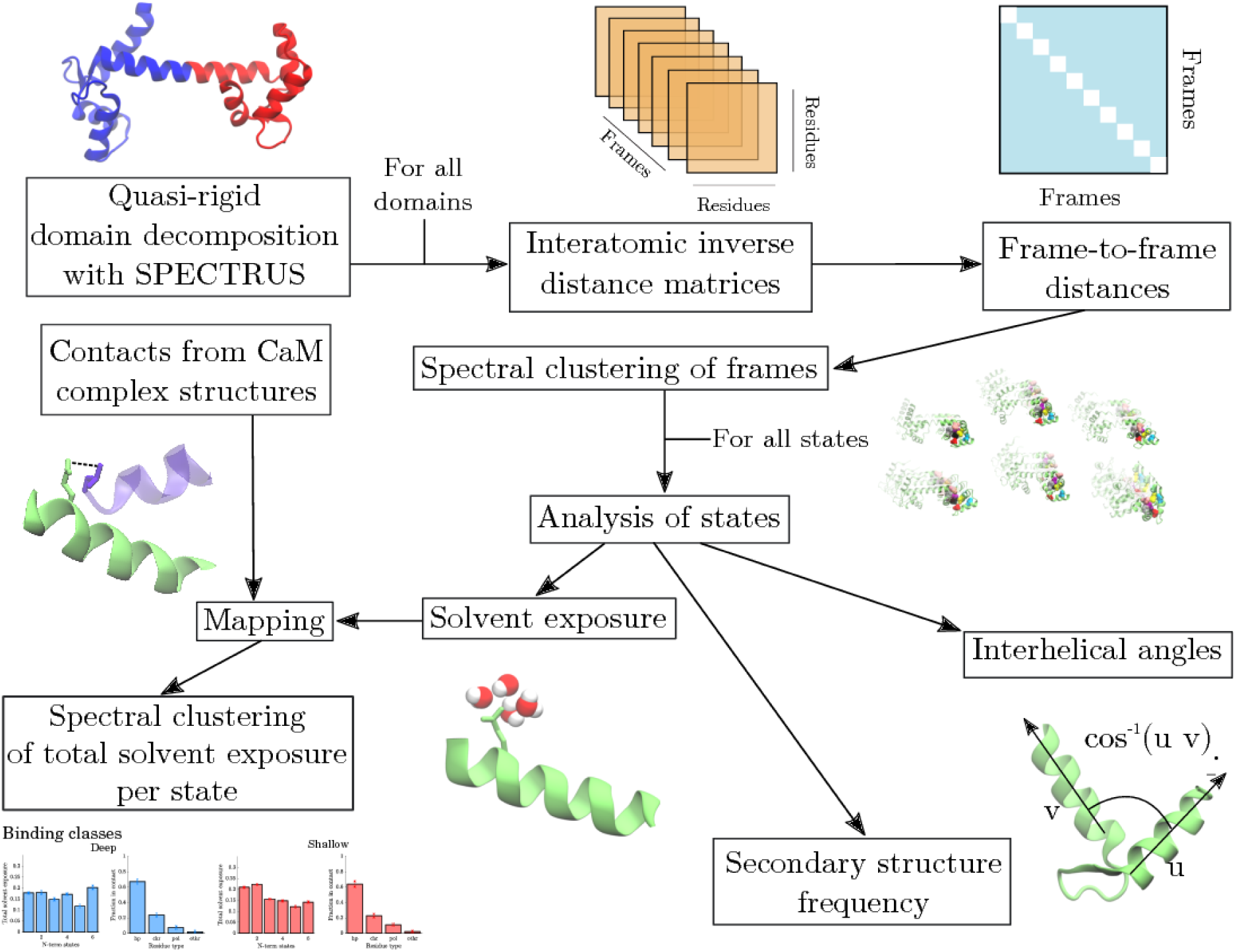
Flow chart of analysis methods. The quasi-rigid domains of CaM were first identified. Then, interatomic inverse distance matrices were used to cluster the frames into states with spectral clustering. The states were analyzed by computing interhelical angles and secondary structure frequencies. Finally, the solvent exposure of each state was mapped to contacts formed in CaM structures, followed by a classification of binding modes with spectral clustering

In a first step, the protein was divided into quasi-rigid domains using SPECTRUS [57]. This procedure exploits fluctuations between residues to determine which parts of the protein move together. The clustering and post-processing described hereafter were done on each quasi-rigid domain.

To further reduce dimensionality and complexity of the dataset while preserving the most important features, we used spectral clustering [58]. The advantage of spectral clustering compared to regular clustering techniques like *k*-means is three-fold. First, it is able to accurately assign points to non-convex clusters. Second, non-linear dimensionality reduction is intrinsic to the algorithm. For high-dimensional data, such as MD trajectories, non-linear dimensionality reduction improves clustering by circumventing the curse of dimensionality where the sparsity of the data increases with increased dimensionality [59]. Third, the number of clusters is the same as the number of dimensions onto which the points are projected. This feature becomes advantageous as the number of free parameters is reduced.

In spectral clustering, the data manifold is first approximated by a graph with adjacency, or similarity, matrix *A*. It contains the local relationships between points and is constructed given the matrix of distances *d*. The distances *d*_*ij*_ are passed through a radial basis function, or Gaussian kernel, with scaling parameter *σ*, yielding the graph edge-weights

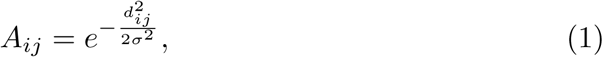

where *d*_*ij*_ is the dissimilarity between conformation *x*_*i*_ and *x*_*j*_. The geodesic distance between two points is the distance between these points along the manifold, the shortest distance on the graph. The size of the scaling parameter, *σ*, influences the accuracy of geodesic distances, and should not be too large nor too small. A too large *σ* would yield short-cuts, thus causing non-convex clusters to be poorly defined. A too small *σ*, on the other hand, would result in a disconnected graph. Here, *σ* is the average nearest neighbor distance between configurations. The random walk matrix, related to the Laplacian [58], is then constructed

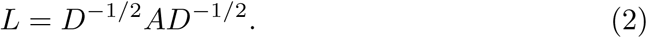

The degree matrix, *D*, is diagonal with *D*_*ii*_ being the degree of node *i*. The first *k* eigenvectors are computed and normalized per row to obtain points projected onto the *k*-sphere. These points are clustered into *k* clusters using *k*-means with centers projected onto the sphere. The choice of number of clusters is guided by the maximum eigengap, the difference between two consecutive eigenvalues. The representative structure of a cluster is chosen as the structure with smallest RMSD with respect to the other structures in the cluster. A cluster with all structures represents the conformational heterogeneity of one state.

Here, in practice, the dissimilarity between conformations, *d*_*ij*_, is measured as the distance between contact maps of inverse inter-atomic distances (iiad-cmap). This general metric is effective for all proteins and, unlike root-mean-square-deviation (RMSD), does not rely on structural alignment. The inverse distances make larger distances small so that far-away motions are cancelled out, without requiring a cutoff. The frame-to-frame distance matrix is compiled by computing the distances between each frame iiad-cmap.

### Analysis of states

Each state, or cluster of frames, was analyzed to provide statistical information about its specific characteristics and molecular features. The features included interhelical angles, secondary structure and importance of states for target protein binding.

### Extracting molecular features of states

A common approach to study calmodulin conformations is to compute its interhelical angles [2, 24]. The interhelical angles in part define the exposed hydrophobic interface and thus selectivity. The angle, *α*, between two helices, ***u*** and ***v***, was determined by their dot product, ***u v*** = ‖***u‖ ‖ v‖*** cos(*α*). We defined a helix vector as the principal eigenvector of the points corresponding to the heavy backbone atoms in sequence direction. The angles between each pair of helices *A– D* (N-CaM and first half of linker) and *E –H* (second half of linker and C-CaM) were calculated, following the helix definitions from Kuboniwa et. al [2]. The helix boundaries are shown in Table 2. We compared the angles found with the method used here to the results in [2, 24], as well as with the experimentally obtained structures of protein-unbound holo and apo CaM, Table 3, and CaM bound to target proteins, Tables 4-5.

**Table 2.** Residue definition of the calmodulin alpha helices.

| Helix ID | Residues in helix |
| --- | --- |
| **A:** | 6-18 (holo: 13-18) |
| **B:** | 29-38 |
| **C:** | 45-54 |
| **D:** | 65-74 |
| **E:** | 83-91 |
| **F:** | 102-111 |
| **G:** | 118-127 |
| **H:** | 139-145 |

DSSP secondary structure assignment [72] frequencies were computed to detect the fraction of time each residue is involved in *α*-helices, *β*-sheets or coils. In order to easily spot differences between the different conformational states, as well as between the four binding states, the difference in secondary structure frequency was compared to a typical holo ensemble. To be able to evaluate the fluctuations in the holo simulations, the typical holo ensemble was represented by one third of the T-REMD trajectory.

A conformational state of CaM was characterized by the average solvent exposure of each residue. Solvent exposure of a residue was calculated as the number of water-oxygens in contact with any heavy atom of the residue. The cutoff was chosen to be 4.5 Å. This was not only done for each state, but also for the complete holo, apo, C-holo and N-holo ensembles, respectively. To characterize allosteric communication between the two lobes, we calculated the residue solvent exposures of C-holo and N-holo compared to holo and apo ensembles. To obtain the relative solvent exposure of a state, the cluster-specific solvent exposure was divided by the average solvent exposure of all states.

Distances and DSSP secondary structure assignment were calculated with MDtraj [73], and molecular visualization was done with VMD [74].

### Kinetic analysis

We calculated the average time taken to transition from one state to another in the MD simulations. The kinetic rates are taken as the inverse of this average time.

### Propensity of states to exhibit conformational selection

We studied the involvement of the different states (clusters) in binding various targets in a conformational selection mechanism by comparing them with CaM structure in complex with targets. This analysis required four steps: 1) defining the CaM residues in contact in the different available CaM-complex structures, 2) computing the relative solvent exposure of each residue for the different sampled states, 3) mapping these two results, and 4) classifying different binding modes. These steps are depicted in the bottom left part of Fig 3.

A contact in CaM-complexes was defined as a CaM-residue atom being within 4.5 Å of a target-protein residue atom. For an ensemble of structures, the residue pair were said to be in contact if they were within 4.5 Å for 70 % of the structures. A contact vector, *m*^(*k*)^, was devised for each complex, *k*, marking CaM contact residues with ones and CaM non-contact residues with zeros,

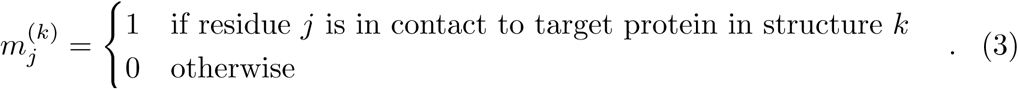

If conformational selection dominates the binding to a target protein, the CaM residues in contact to this protein should be exposed to solvent when CaM is unbound. Therefore, the relative solvent exposure, 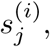 of residue *j* in state *i*, was calculated for each residue. The matching score, or total solvent exposure, for a state *i* and complex *k* was computed as the sum over all residues in contact to the target protein,

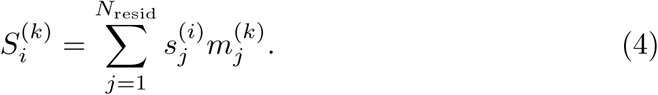

This was normalized over states to provide a distribution for each CaM-complex.

To identify patterns of different classes of binding modes, these distributions were clustered. For this, spectral clustering was performed again, taking the distance matrix *d* to be the distance between the distributions. For each class, the mean distribution, with standard deviation, of total relative solvent exposure was computed. Moreover, the residue types (polar, apolar, charged) involved in the contacts were identified and the average distribution of residue types as well as the standard deviation were calculated.

## Results and Discussion

To characterize the ability of calmodulin to bind to calcium and target peptides through conformational selection, the protocol outlined in Fig 3 was used. The configurations of quasi-rigid domains in molecular trajectories were partitioned with spectral clustering and the obtained conformational states (hereby referred to as states) were analyzed.

### Conformational selection aspects of Ca^*2+*^-binding in the two lobes

We sampled conformations of apo and holo CaM using MD, T-REMD and REST, Table 1. The analysis began by identifying two quasi-rigid domains, namely N-CaM and C-CaM, with SPECTRUS [57]. The identified domains are shown as red and blue CaM ribbons in Fig 3. The clustering and post-processing were therefore carried out on these two domains separately. S1 Fig-S4 Fig show the representative structures of holo and apo CaM obtained after clustering on C-CaM and N-CaM. Together with these states, experimental structures of similar conformations are shown for comparison.

The molecular features were extracted from the conformational states by computing the interhelical angles and secondary structures. Fig 4 shows the interhelical angles of each state (cluster) from the holo simulations, while Fig 5 shows the interhelical angles for each state of the apo simulations. Each dot is a projection of a conformation in interhelical angle space, colored according to the state (cluster) it belongs to. On top of this, experimentally obtained structures are plotted as squares. The white squares denote structures where CaM is not bound to target peptides, Table 3, while pink squares correspond to structures with CaM bound to a target peptide, Table 4 and 5. The black circles are mean values of CaM interhelical angles inferred from NMR data [2, 24], used for comparison. Furthermore, to allow a comparison between interhelical angles in the holo and apo ensemble, the sampled apo conformations are shown as a gray shadow in Fig 4 (holo) and the sampled holo conformations are shown as a gray shadow in Fig 5 (apo). A simple sanity check, S5 Fig-S10 Fig, showed that most states were sampled by all three simulation methods, with a few exceptions. Examples of such are states that are separated by a high free energy barrier and therefore not sampled by the regular MD simulations within these timescales, or states that could not be sampled by the replica exchange simulations because of the relatively short time scales simulated. Similar conclusions can be drawn from the kinetic analysis of MD trajectories (S11 Fig).

**Table 3.** Experimental structures of calmodulin used to compare interhelical angles with the MD dataset.

| **PDB ID** holo | **PDB ID** apo |
| --- | --- |
| 2K0E [60] | 1LKJ [36] |
| 1CLM [61] | 1DMO [24] |
| 1CLL [62] | 1CFC [2] |
| 1EXR [63] |  |
| 1OOJ [64] |  |
| 1OSA [65] |  |
| 1PRW [66] |  |
| 1UP5 [67] |  |
| 3CLN [35] |  |
| 4CLN [68] |  |
| 4BW7 [69] |  |
| 4BW8 [69] |  |
| 3IFK [70] |  |
| 1AK8 [71] |  |

**Table 4.**
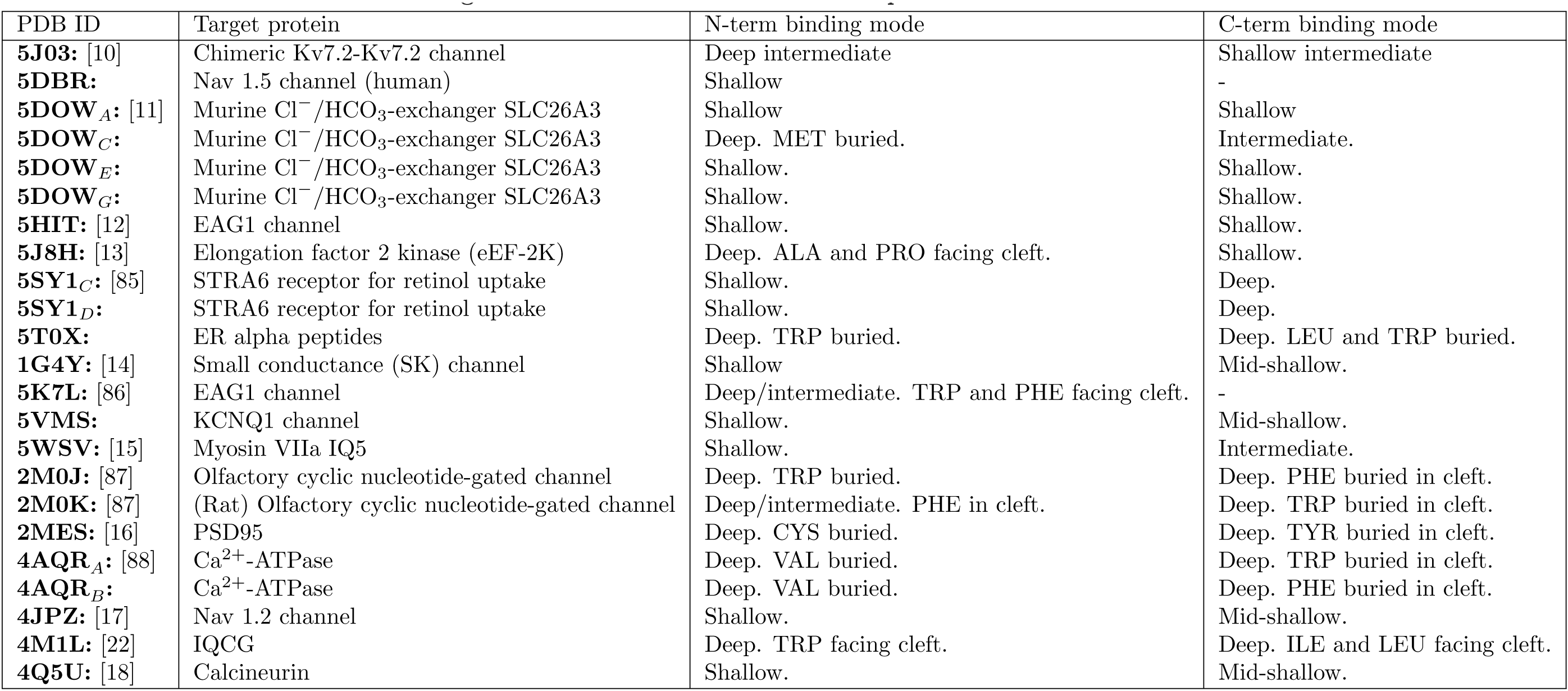
Structure PDB ID and binding mode identified for holo CaM complexes.

**Table 5.**
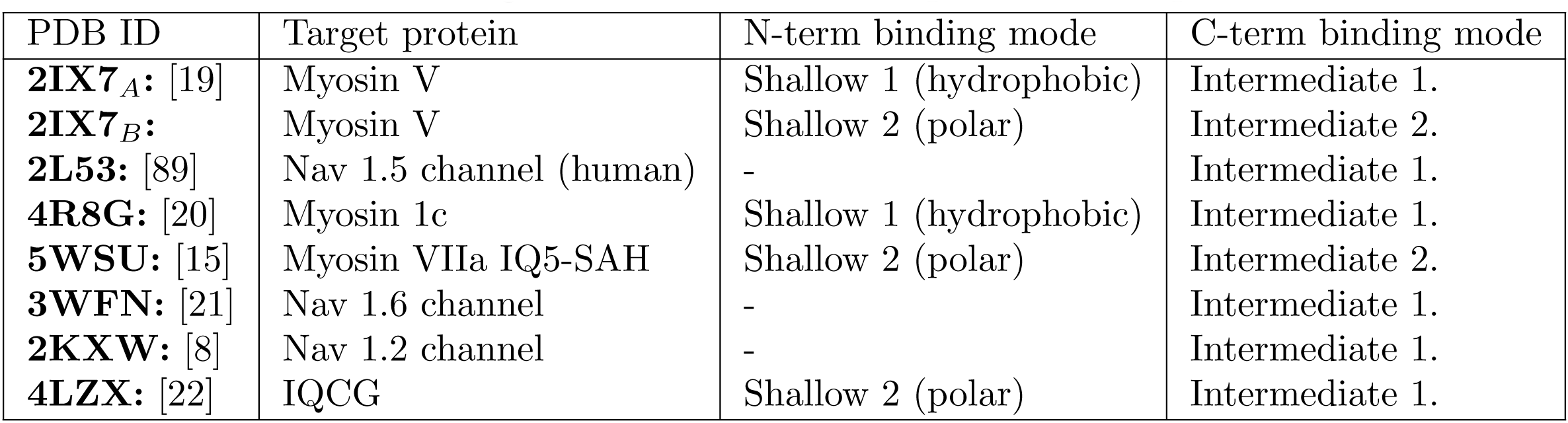
Structure PDB ID and binding mode identified for apo CaM complexes.

| PDB ID | Target protein | N-term binding mode | C-term binding mode |
| --- | --- | --- | --- |
| **2IX7<sub>A</sub>**: [19] | Myosin V | Shallow 1 (hydrophobic) | Intermediate 1. |
| **2IX7<sub>B</sub>**: | Myosin V | Shallow 2 (polar) | Intermediate 2. |
| **2L53**: [89] | Nav 1.5 channel (human) | - | Intermediate 1. |
| **4R8G**: [20] | Myosin 1c | Shallow 1 (hydrophobic) | Intermediate 1. |
| **5WSU**: [15] | Myosin VIIa IQ5-SAH | Shallow 2 (polar) | Intermediate 2. |
| **3WFN**: [21] | Nav 1.6 channel | - | Intermediate 1. |
| **2KXW**: [8] | Nav 1.2 channel | - | Intermediate 1. |
| **4LZX**: [22] | IQCG | Shallow 2 (polar) | Intermediate 1. |

**Fig 4.**
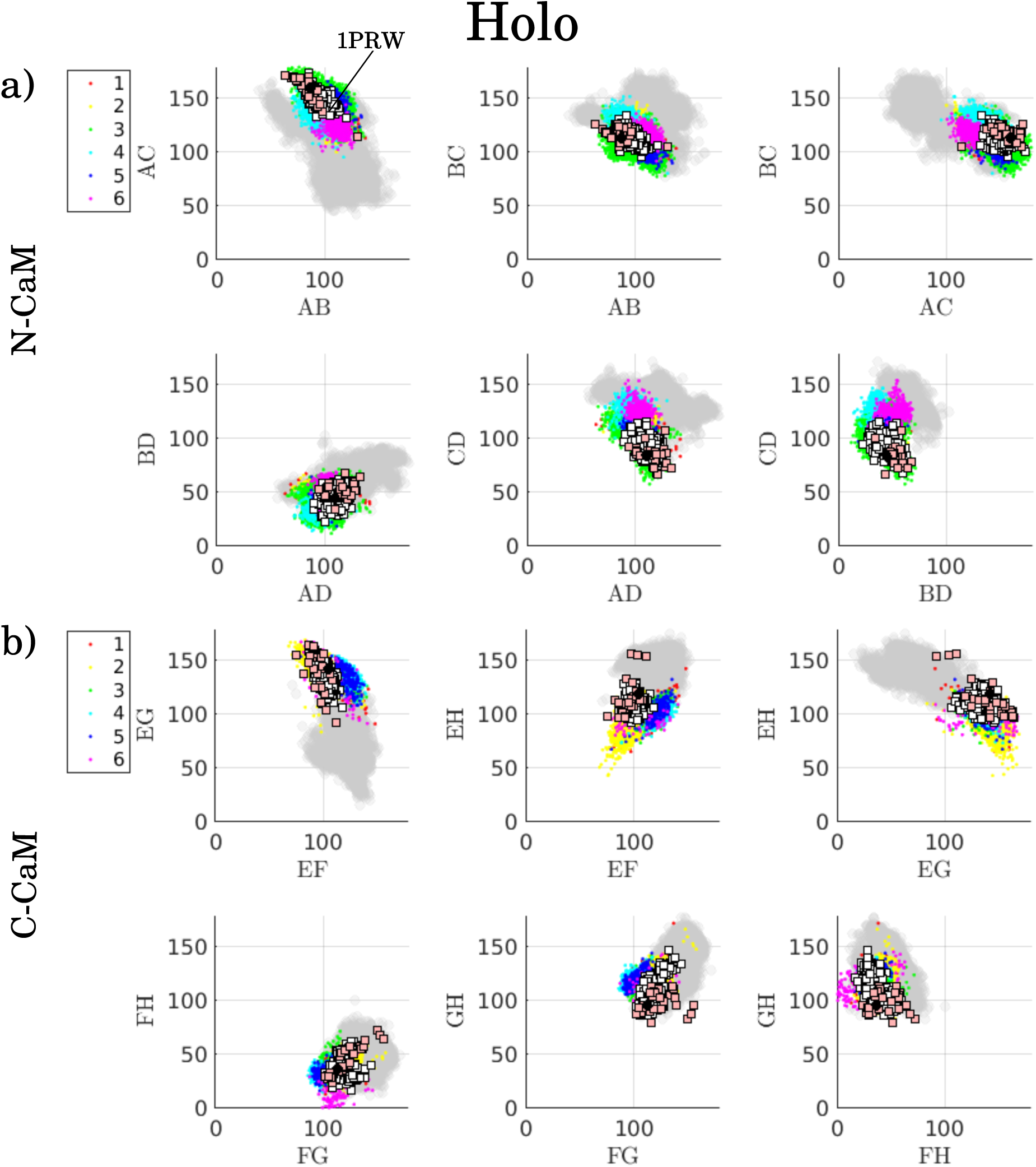
a) N-CaM and b) C-CaM states (clusters) projected onto interhelical angles of holo CaM, colored by their respective assigned cluster. The gray dots correspond to the apo ensemble. The white squares are experimentally determined structures of protein-unbound holo CaM, while pink squares are experimentally determined structures of protein-bound holo CaM. The black circles respresent values of CaM interhelical angles inferred from NMR data [2, 24].

**Fig 5.**
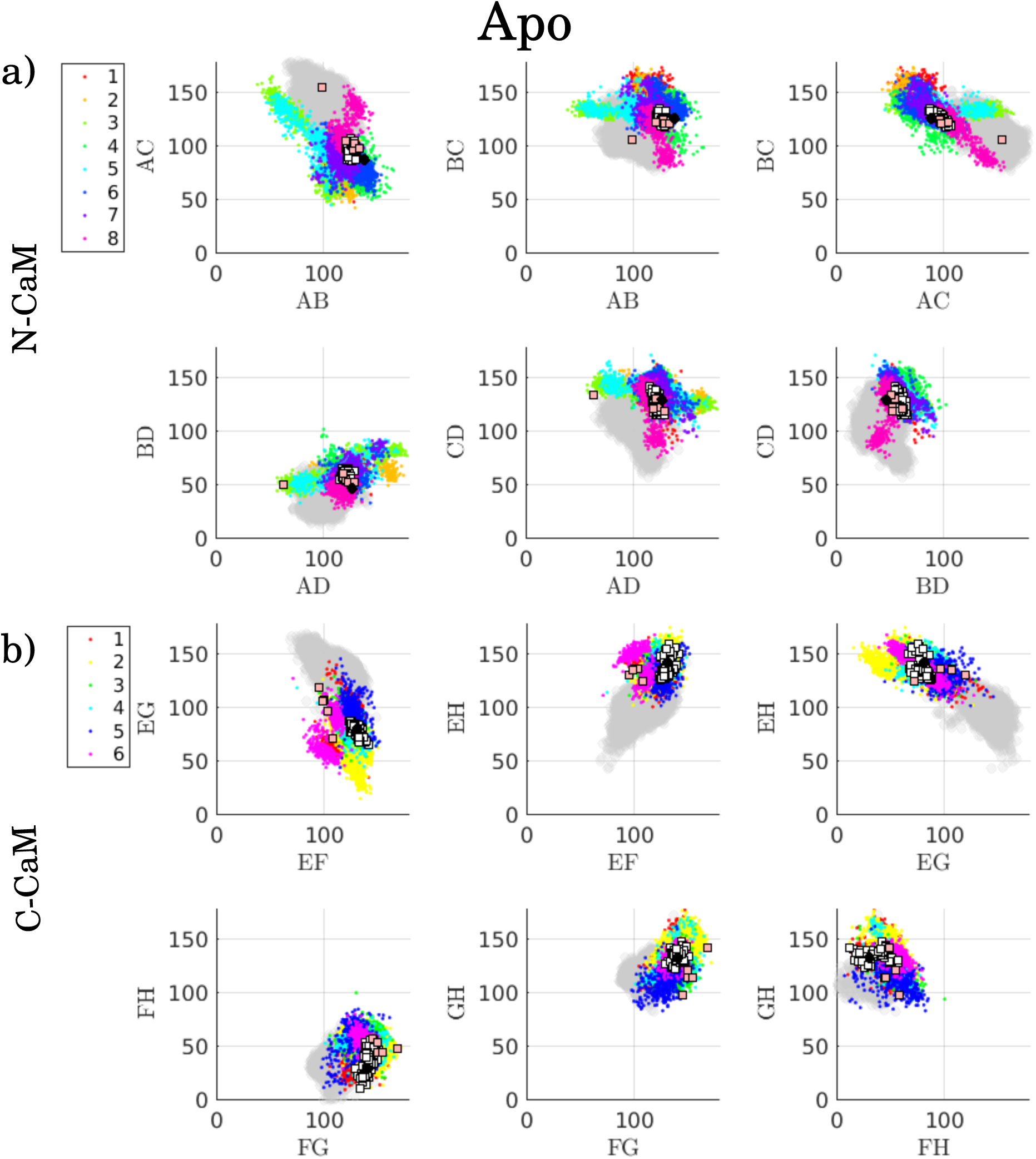
a) N-CaM and b) C-CaM states (clusters) projected onto interhelical angles of apo CaM, colored by their respective assigned cluster. The gray dots correspond to the holo ensemble. The white squares are experimentally determined structures of protein-unbound apo CaM, while pink squares are experimentally determined structures of protein-bound apo CaM. The black circles represent values of CaM interhelical angles inferred from NMR data [2, 24].

A change in interhelical angles due to Ca^2+^-binding can be seen in Figs 4 (holo) and 5 (apo), in agreement with previous work [2, 24, 26] (see S1 Text for more extensive description and details). We note also that the conformational ensemble generated by MD simulations is broader than the ones observed experimentally for both protein-bound and unbound CaM. In agreement with NMR studies of holo CaM [60], the protein-unbound CaM adopts the conformations of CaM bound to specific target proteins, which is seen as an overlap between the pink squares and the MD data (colored dots) in Fig 4, thus hinting at a possible conformational selection mechanism of target protein binding.

Fig 6 displays the difference in secondary structure frequency for each residue compared to holo. Red dots denote the helical, black the strand and blue the coil content. Positive values indicate a gain in secondary structure element compared to holo, and negative ones a loss. The helical break around residue 74-82 [26–28] is seen to different extents in all Ca^2+^-binding states of CaM, Fig 6. This break occurs in the linker and yields a compact state with the two lobes in contact. This compact state resembles the binding mode where CaM is wrapped around the target protein and was suggested to be an intermediate conformation during target protein- and Ca^2+^-binding [69, 75]. Furthermore, binding Ca^2+^ rearranges the helices in the lobes and pins the beta sheets between residues 27/63 and 100/136 instead of 28/62 and 99/137, Fig 6. Apo CaM is more flexible, allowing the beta sheets to shift between 26-28/62-64 and 99-101/135- 137, Fig 6. This highlights the rigidity of holo compared to apo CaM. Due to the flexibility of apo, a C-CaM state (Fig 5.b, state 2) is found where the end of the fourth binding loop is involved in a beta sheet, deforming the binding loop, in agreement with [31]. We hypothesize that this state may inhibit Ca^2+^-binding. In this state, residues 129-131 are more prone to form a beta sheet with 99-101 and 135-137, S12 Fig. This extra sheet does not form in apo N-CaM because the coiled loop between helices C and D is slightly shorter than the one in apo C-CaM between helices G and H. The flexibility obtained by removing Ca^2+^ from C-CaM allows these beta sheet formations to occur.

**Fig 6.**
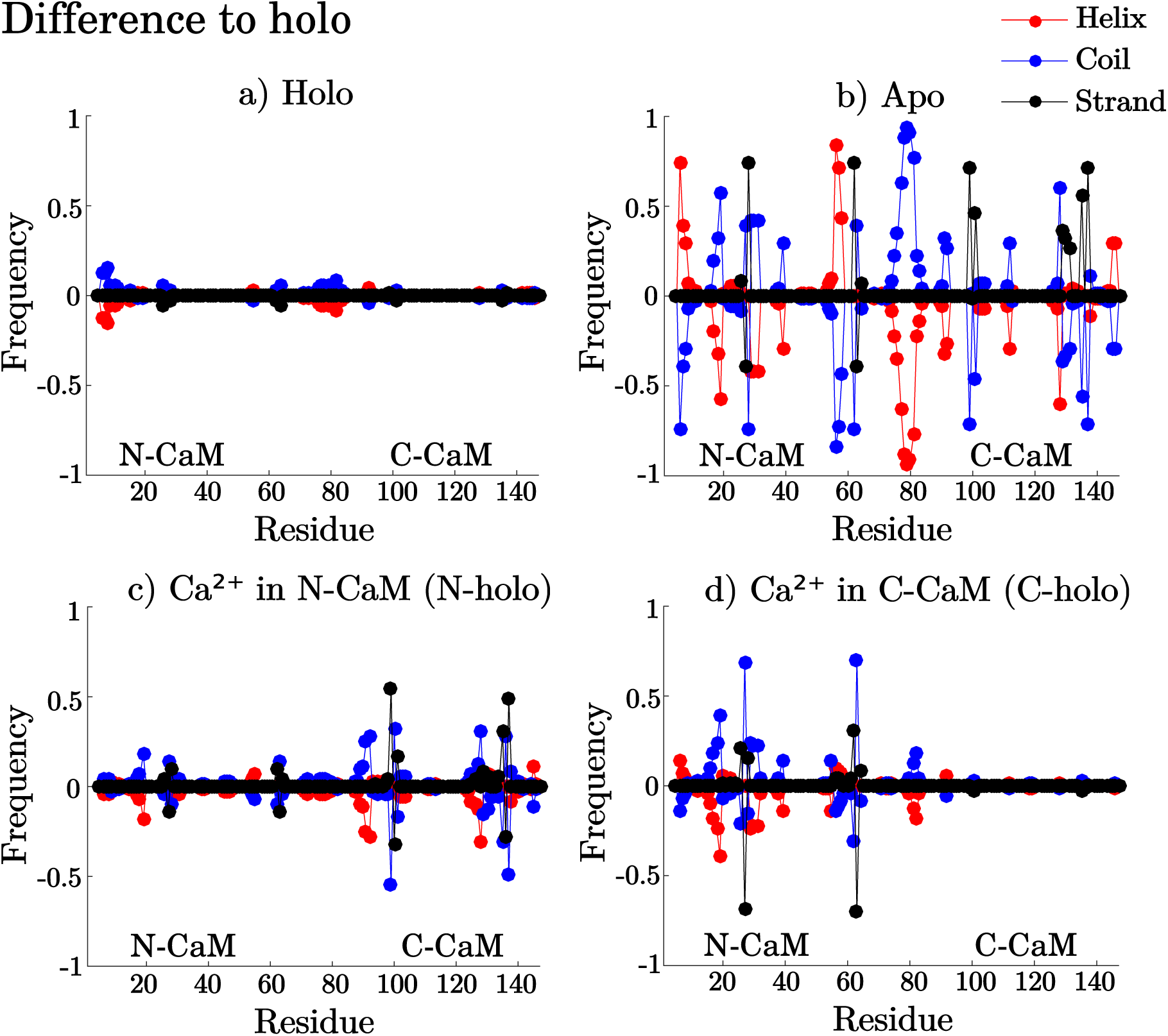
Secondary structure frequency difference to holo for a) complete holo,b) C-holo, c) N-holo and d) apo ensembles. Helix frequency is shown in red, strand in black and coil in blue. Above zero denotes gained secondary structure compared to holo while below shows the loss in secondary structure frequency

To investigate the possibility for conformational selection during the Ca^2+^- binding process, we searched for potential overlaps in interhelical angle space between apo and holo. We observe holo-like states in apo N-CaM, Fig 5.a (states 3, 5 and 8, plotted with colors lime, cyan and cerise), as well as C- CaM holo-like states that have been reported in previous work [31, 76], Fig 5.b (states 1 and 5, depicted as red and blue). These states could potentially aid Ca^2+^-binding though conformational selection. The holo N-CaM states 4 and 6 (shown by cyan and magenta dots) overlap with the apo ensemble, Fig 4.a. Their secondary structure frequencies are plotted in S13 Fig, which show similar secondary structure frequencies as apo (Fig 6.b), with a shift of beta sheet to residues 28/62. The C-CaM interhelical angles of holo and apo overlap more than the N-CaM interhelical angles do. The overlap in interhelical angles and shift of beta sheet in N-CaM, make these states likely intermediate states in Ca^2+^-binding, involved in conformational selective binding of Ca^2+^.

To assess whether Ca^2+^-binding allosterically modulates the conformational ensemble of the other lobe, we used the MD, T-REMD and REST datasets of C-holo (Ca^2+^ in C-CaM) and N-holo (Ca^2+^ in N-CaM), Table 1. Solvent exposure for each residue in N-holo and C-holo was compared to the apo (Fig 7.a,c), and holo (Fig 7.b,d) ensembles. This shows that the Ca^2+^-free lobe tends to transition to an apo-like conformational ensemble, while the Ca^2+^-bound lobe stays in the holo-like ensemble. Although there is a small allosteric influence between the lobes, as seen in Fig 6.c,d, where N-CaM mostly lacks beta sheet in C-holo, the solvent exposure analysis indicates that the overall conformation of one lobe is mostly independent of the conformation of the other. These results indicate that binding of Ca^2+^ to one of the lobes does not yield a significant population shift in the other lobe and thus that binding is likely not conformationally cooperative between the two lobes. Cooperative binding between sites and lobes has been investigated but the results obtained in different studies have been largely inconsistent, thus making it difficult to validate our results. This is because many different parameter sets in the models used to fit the experimental data perform equally well [77].

**Fig 7.**
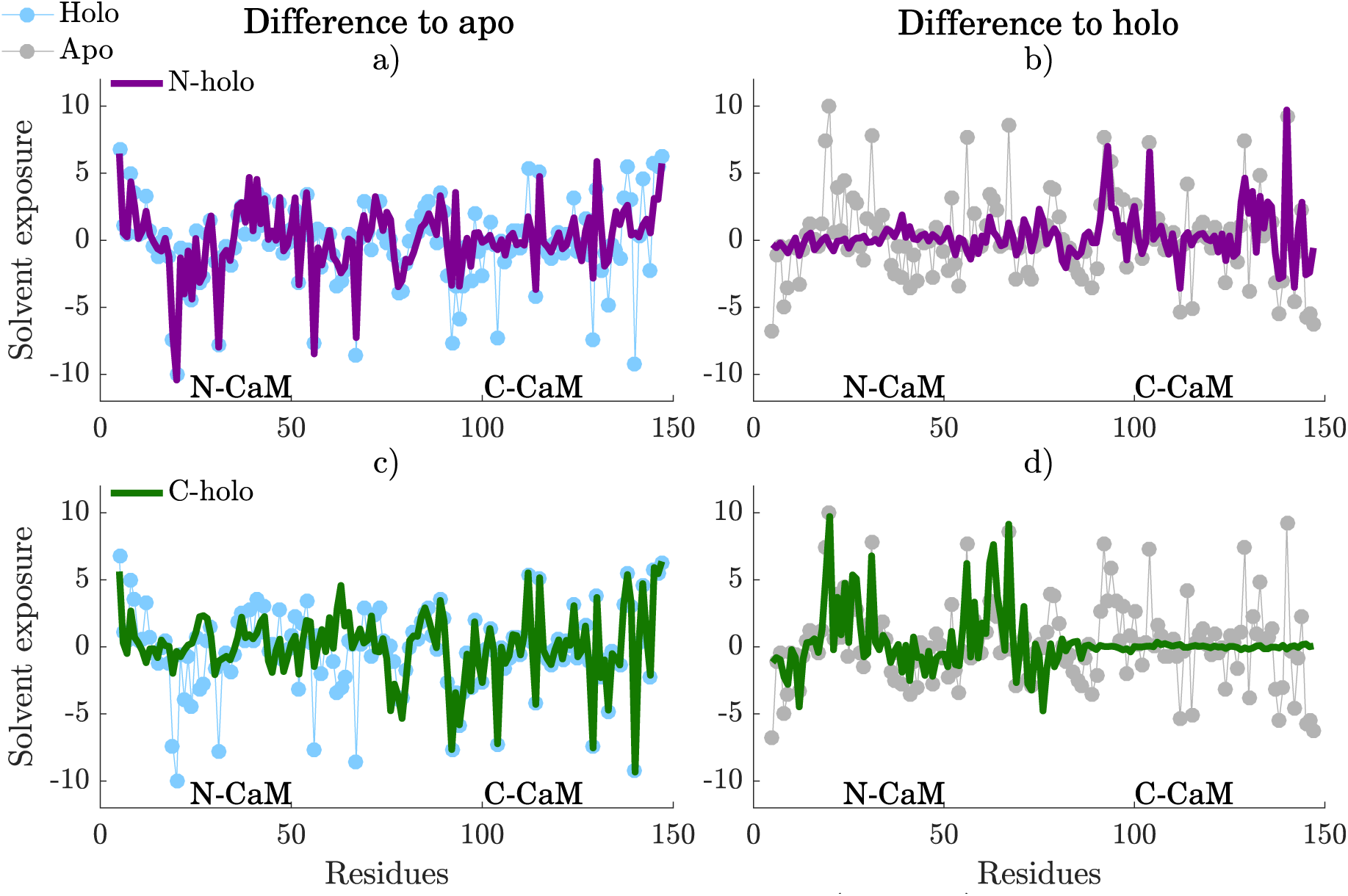
Solvent exposure difference of N-holo (purple) and C-holo (green) to a, c) apo and b,d) holo. The difference in solvent exposure to holo of both N-holo and C-holo are compared to the solvent exposure difference of apo (gray) to holo, while the difference in solvent exposure to apo of the two are compared to the solvent exposure difference of holo (blue) relative to apo.

### Conformational selection aspects of target protein-binding

To investigate conformational selection aspects of calmodulin binding to target proteins, the relative solvent exposure of the different CaM residues in different states was calculated and mapped to the contacts in a set of CaM complexes, Tables 4 and 5 and Fig 3. The underlying idea is that in a conformational selec-tion mechanism, water molecules around an exposed residue will be replaced by the target peptide. Therefore, the residues that are in contact with the target peptides should be found exposed to solvent in a state involved in a binding mechanism dominated by conformational selection. To then characterize different modes of binding, the distributions of total solvent exposure per state were clustered and divided into classes, as described in the Methods section.

### Holo C-CaM complexes can be divided into five classes according to binding depth

The clustering of distributions of total relative solvent exposure of residues involved in binding holo C-CaM complexes (Table 4) yielded five subclasses. Through visual inspection, we aligned these along a depth gradient with deep, intermediate and shallow subclasses, Fig 8, S14 Fig and S15 Fig. Deep binding is characterized by a large hydrophobic target peptide residue buried in the CaM hydrophobic cleft. It is favored by states 3 and, especially, 6. These states expose hydrophobic residues from deep within the cleft simultaneously; PHE92, LEU105, ILE100, VAL121, VAL136 and PHE141. State 6 is characterized by parallel F and G helices, Fig 4.b, and lacking beta sheet structure, S16 Fig.

**Fig 8.**
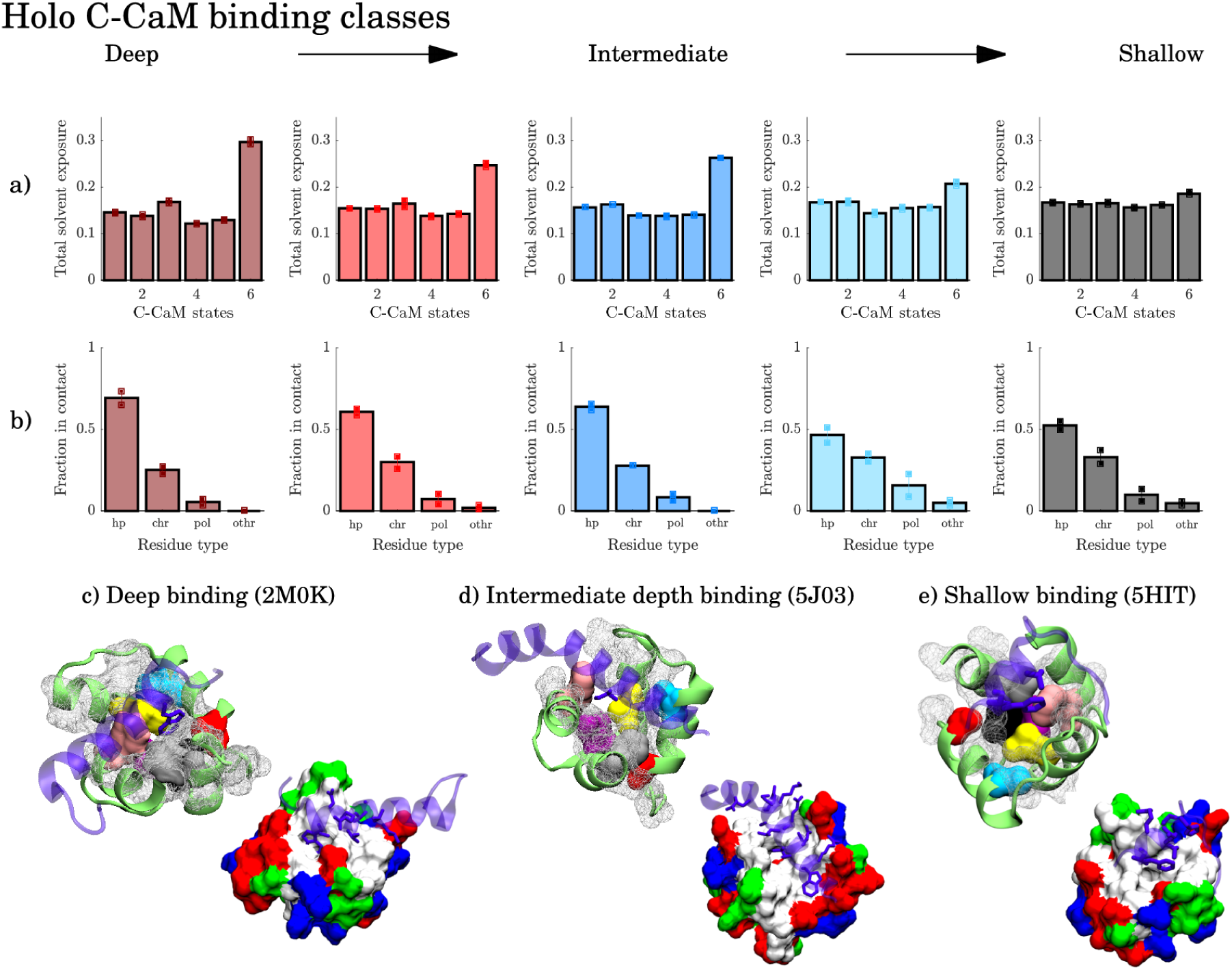
Representative models from the three binding classes of holo C-CaM with deep binding in c) olfactory cyclic nucleotide-gated ion channel (OLFp), intermediate binding in d) chimeric Kv7.2 - Kv7.3 ion channel and shallow binding in e) EAG1 ion channel. Color-highlighted residues are hydrophobic residues showing distinct signals in the contact-solvent exposure analysis. The other hydrophobic residues are shown by the white mesh.Associated to each class is a) the average total solvent exposure distribution per state and b) the average distribution of contact types formed in the CaM-complexes (lower row). The standard deviation is indicated on the bars.

This state, which could be favorable for binding target proteins, is a high free energy state, only observed in the temperature enhanced simulations, S17 Fig. Binding is also likely when CaM is in the state 3 conformation, but binding to state 3 may require more subsequent rearrangements than binding to state 6. An example of deep binding is shown by the contact interface of CaM-CNG ligated ion channel (2M0K) where the a tryptophan is buried in the hydrophobic cleft, Fig 8.c. Coincidentally, the residues that give rise to significant signals in relative solvent exposure, S18 Fig-S20 Fig and S1 Fig, are also highly conserved in eukaryotes through evolutionary time [78].

Intermediate depth binding specifically favors state 6 and 2, Fig 8. In intermediate depth binding, the residue pointing towards the cleft is smaller than in deep binding, as in the case of CaM-Kv7.2-Kv7.3 (5J03) binding interface with a leucine pointing towards the cleft, Fig 8.d.

In the superficial binding class, the residues in contact are roughly equally exposed to solvent in all states, Fig 8.a and S18 Fig-S20 Fig. As seen in the distribution of residue types that are part of contacts, Fig 8.b, less hydrophobic contacts are formed for more shallow binding modes. Moreover, the shallow binding modes include a larger fraction of charged residues, and do not favor any particular state since the residues needed for contact are always equally exposed to solvent. Thus, the long-ranged electrostatic interactions of the charged residues may initiate binding, yielding fast and promiscuous binding, while the CaM hydrophobic pocket of deeper binding may account for selectivity and be involved in binding mechanisms dominated by conformational selection.

### Holo N-CaM complexes can be divided into two classes according to binding depth

For holo N-CaM binding, clustering enabled to identify 2 classes corresponding to deep binding and shallow binding, Fig 9, S21 Fig and S22 Fig. In shallow binding, states 1 and 2 appear more favorable than the rest. The key feature of shallow binding resides in smaller hydrophobic target protein residues facing the CaM hydrophobic cleft, Fig 9.b.

**Fig 9.**
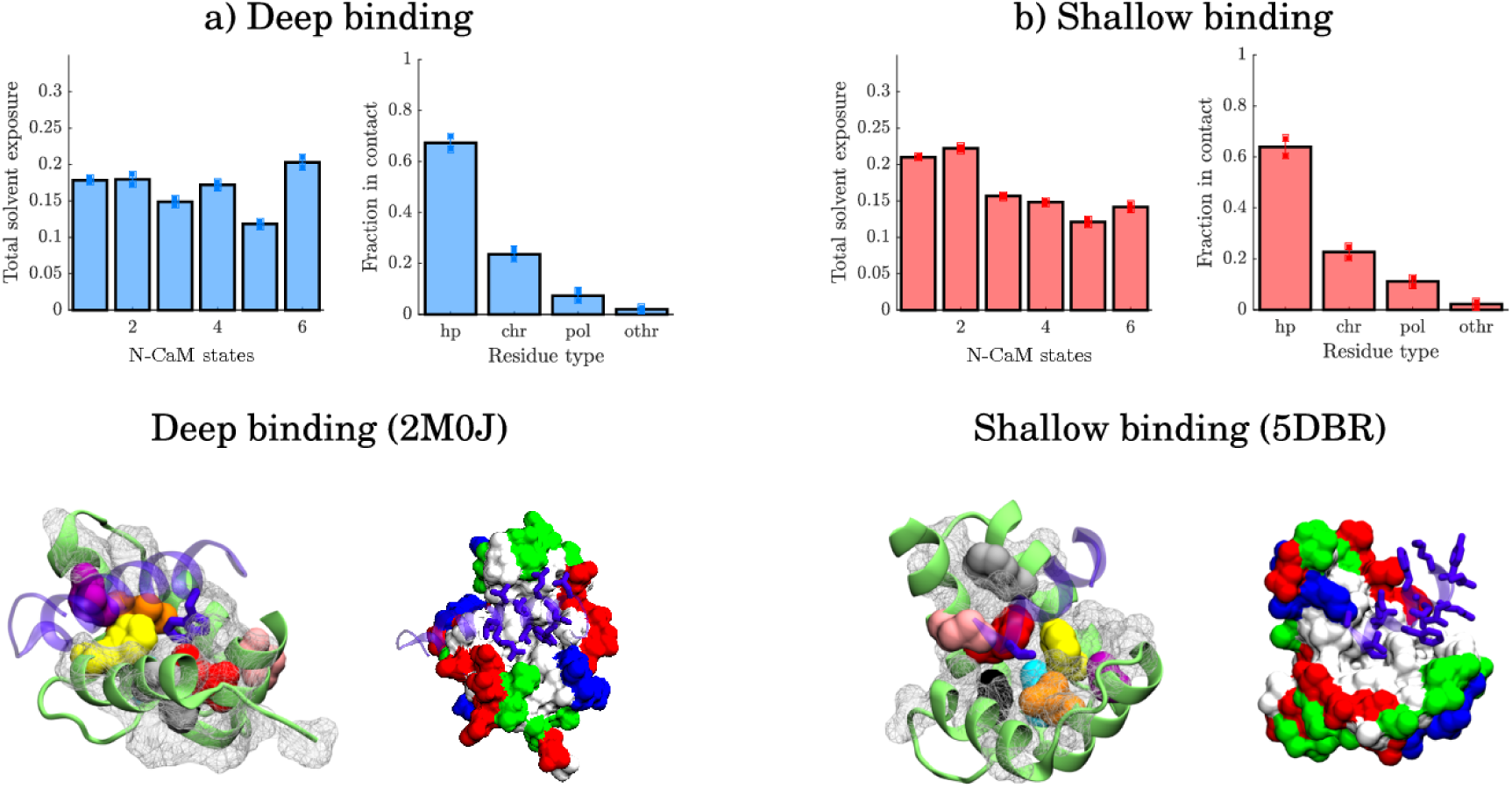
Representative models from the two binding classes of holo N-CaM with deep binding in a) olfactory cyclic nucleotide-gated ion channel (OLFp) and shallow binding in b) (human cardiac) Nav1.5 ion channel.Color-highlighted residues are hydrophobic residues showing distinct signals in the contact-solvent exposure analysis. The other hydrophobic residues are shown by the white mesh. Associated to each class is the average total solvent exposure distribution per state (left) and the average distribution of contact types formed in the CaM-complexes (right). The standard deviation is indicated on the bars.

For deeper binding, a mix of states gives exposure to all necessary contact residues, namely states 1, 2, 4 and 6, Fig 9.a. State 6 is slightly more favorable for complexes with deeper binding, associated with more hydrophobic contacts S21 Fig. However, since not one state exposes all necessary residues, the binding of the N-CaM could be dominated by induced fit. After binding to any state, the rest of the residues necessary for binding will be exposed to reduce free energy. In state 4, CaM binds its own helix A, S2 Fig. This state has been observed before with N-CaM forming salt bridges to the linker [79]. It is also represented by structure 1PRW [66] which overlaps state 4 in the interhelical angle space, Fig 4.a. It may display a possible intermediate conformation during a flexible binding process in the N-term.

The hydrophobic cleft of N-CaM is not as open as in C-CaM. This is reason to believe that while C-CaM binding may be dominated by conformational selection, N-CaM binding may need a more flexible mechanism with intermediate states to facilitate deep binding. Also here, however, deep binding involves a large hydrophobic target protein residue buried in the CaM cleft, Fig 9.a.

As in C-CaM binding, the depth of binding in the N-CaM hydrophobic cleft determines which interacting residues and therefore which states are more favorable for binding. The contacts in deeper binding are formed through a combination of CaM residues ILE27, LEU32, VAL35, ILE63, PHE68 and MET71, S23 Fig-S25 Fig and S2 Fig. These contacts are here associated to a large hydrophobic residue from the target peptide being buried or facing the hydrophobic cleft. Furthermore, these CaM residues are also reported as highly conserved in eukaryotes. [78]

In summary, states 1 and 2 expose residue PHE68 and MET71 which are required even in shallow binding. Therefore, state 1 and 2 are presumably favorable for initiating binding to both shallow and deeper binding modes. State 6 exposes residues in contact in slightly deeper binding and state 4 is probably an intermediate state visited in on the way towards deep binding.

### Apo CaM target-protein binding is more shallow than holo CaM binding

Fig 10.a and S26 Fig show the classification of apo C-CaM binding. In this case the classes are divided according to whether contacts are formed with CaM residue LEU105 or PHE141 (in which case state 5 is favored) or with PHE89 (in which case state 6 is favored), Fig 10, Table 5 and S27 Fig. These residues are more exposed to solvent in the states that favor binding to the analyzed target proteins, suggesting that they are important for C-CaM binding.

**Fig 10.**
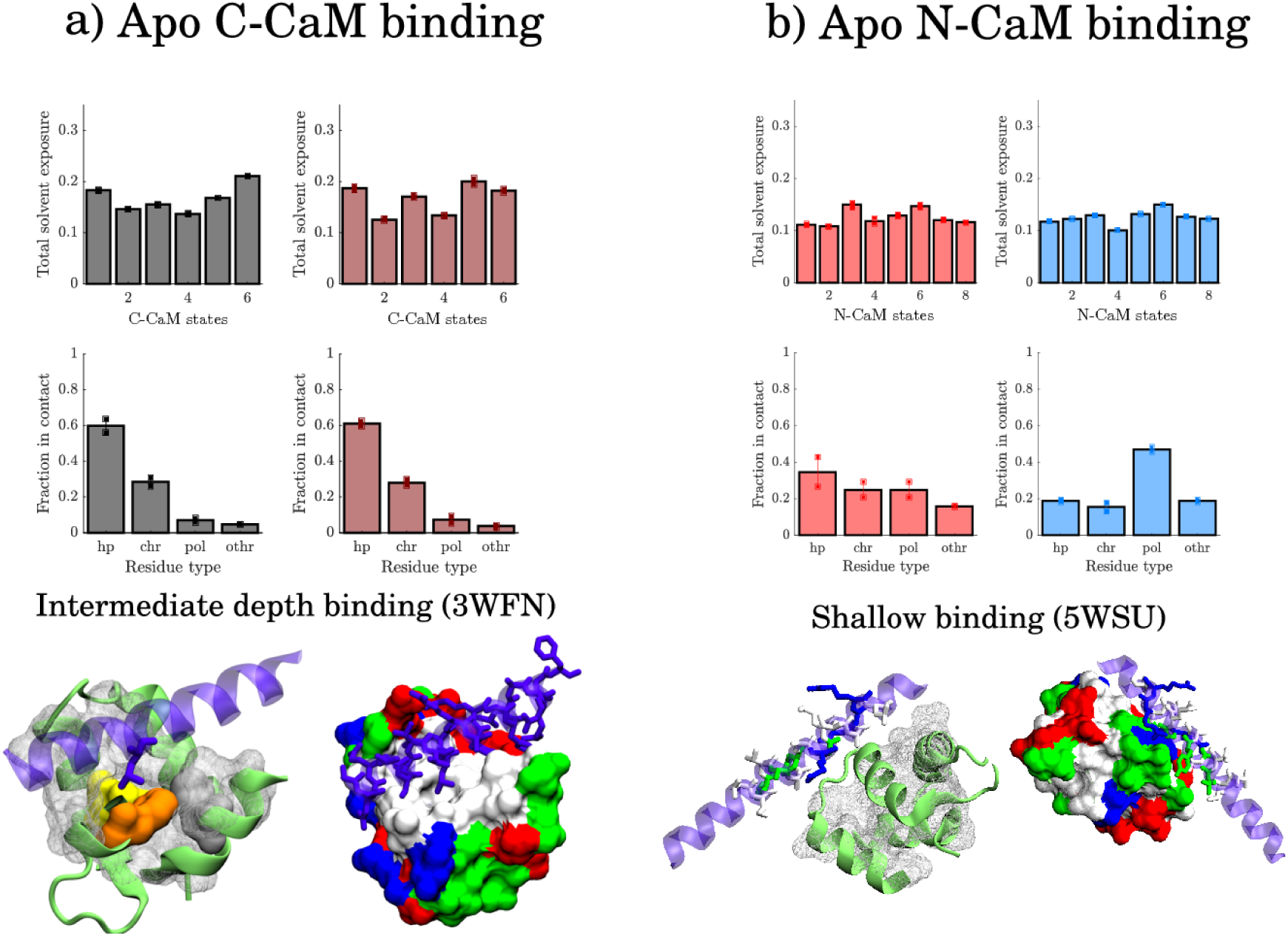
Representative models from the binding classes of apo C-CaM and N-CaM with C-CaM intermediate binding to a) Na_V_1.6 ion channel and N-CaM shallow binding to b) Myosin VIIa. Color-highlighted residues are hydrophobic residues showing distinct signals in thecontact-solvent exposure analysis. The other hydrophobic residues are shown by the white mesh. Associated to each class is the average total solvent exposure distribution per state (upper row) and the average distribution of contact types formed in the CaM-complexes (lower row). The standard deviation is indicated on the bars.

Interestingly, mutating CaM either at PHE89 and PHE141 to LEU causes arrhythmias and long QT syndrome [80–82]. Moreover the Ca^2+^ affinity has been shown to decrease for the PHE141 to LEU mutation [83]. This residue is, in fact, more exposed in state 1 and 5 (S27 Fig and S3 Fig), the two previously identified holo-like states, Fig 5, which indicates that this residue is important for both calcium as well as target protein binding.

As in holo, apo C-CaM target protein binding involves a larger fraction of hydrophobic residues than apo N-CaM binding. When binding Na_V_1.6 (3WFN), for example, a LEU is buried in the CaM hydrophobic cleft, Fig 10.a. In the structures of apo N-CaM binding, however, we only observe superficial binding, Fig 10.b and S28 Fig. An example is binding to Myosin VIIa (5WSU), where a VAL is facing the cleft, Fig 10.b. The two classes from clustering instead distinguish whether polar contacts or more hydrophobic contacts are formed. The hydrophobic residues are well packed within N-CaM and a lack of hydrophobic cleft characterizes shallow binding.

### Comparison of apo and holo target peptide binding

For apo CaM binding, the residues that bind to target proteins are in general equally exposed to solvent in all states, S27 Fig and S29 Fig. Therefore, apo does not need to stay in distinct states for binding, but uses its configurational flexibility to adjust the binding pose after initial contacts are made. This flexibility that stems from the low free energy barriers and diffusivity of apo CaM [43], is possibly linked to apo not showing a diverse set of binding interfaces. It has previously been proposed that C-CaM in N-holo is semi-open [25], which would facilitate binding to target proteins, but later it was shown that the semi-open state exists in even pure apo C-CaM [2, 24, 31] and apo C-CaM complexes [8]. Here, the semi-open state is shown to exist in the unbound apo C-CaM ensemble, whereas the N-CaM ensemble is mostly closed. This is in line with the finding where the apo C-CaM interhelical angles overlap holo more than the interhelical angles of apo N-CaM, Fig 5.

Despite the overlap of interhelical angles, and thus semi-openness of apo C- CaM, the true open state and deep binding with a large hydrophobic residue buried in the CaM hydrophobic pocket is an effect of Ca^2+^ and thus only observed in holo CaM. Binding to the protein is then most likely facilitated by the holo C-CaM cluster 6, where the loss of beta sheet opens the cleft and exposes a larger hydrophobic pocket, allowing for deeper binding, S16 Fig, S1 Fig and These well-defined states of holo allow for binding to target peptides in a mechanism dominated by conformational selection.

## Conclusions

We used molecular dynamics and temperature enhanced MD with an agnostic spectral clustering scheme to search for conformational selection of CaM Ca^2+^ and target protein binding. We found that the Ca^2+^-state of one lobe does not significantly influence the conformation of the other lobe, but binding Ca^2+^ may occur through conformational selection facilitated by a transition state that is visited by both apo and holo CaM. It is observed here by overlapping conformations in interhelical angle space. In N-CaM, the transition between apo and holo likely involves a beta sheet residue shift.

The target protein binding modes of apo CaM are more shallow than those of holo CaM, and the charged interactions dominate due to the induced burial of the hydrophobic binding interface from Ca^2+^-depletion. The residues involved in binding are equally exposed in apo CaM states and thus all observed states are equally likely to initiate binding to the target protein.

The notion that not only the linker, but also the binding interface of C-CaM shows configurational flexibility has previously been proposed both through extensive MD simulations [31], but also in structural studies [2, 24, 25]. The conformation of C-CaM interface is suggested to vary more than the N-CaM interface [25]. Here, holo C-CaM shows distinct states exposing hydrophobic residues that are otherwise screened from water. Such states are absent in N-CaM, which also exhibits less binding heterogeneity than C-CaM in the CaM-complex struc-tures. This is seen as the shallow class of C-CaM show more shallow binding than the shallow class of N-CaM, while the deep binding class of C-CaM shows deeper binding than the deep binding class of N-CaM. The distinct states of holo C-CaM with a clear mapping to bound structures indicate a tendency for C- CaM binding to be selective and dominated by conformational selection, while N-CaM, which lacks this clear mapping, likely binds through a more flexible mechanism involving intermediate states.

For general protein-ligand binding, strong and long-ranged interactions have been thought to favor binding dominated by induced fit, while short-ranged interactions would favor conformational selection. [84] Our results support and extend this idea to CaM by showing that weak hydrophobic interactions dominate deep binding modes which indeed tend to be dominated by conformational selection in C-CaM. In the flexible binding of N-CaM, on the other hand, hydrophobic interactions occur less frequently. We hypothesize that the long-ranged electrostatic interactions of the N-CaM charged residues may initiate fast binding while the hydrophobic pocket in C-CaM may account for selectivity. Previously published studies using NMR [23] proposed that C-CaM is selective while N-CaM binds afterwards through induced fit mechanism, or a coupled conformational selection mechanism initiated by C-CaM [60]. This is in line with the argument given here on C-CaM selectivity and N-CaM flexibility.

This study opens the way towards understanding the process and mechanisms behind calmodulin Ca^2+^-sensing. In a next step, the second aspect of binding, induced fit, may be studied. The full mechanism of specific target peptide binding could also be simulated starting from the conformational selection state of CaM identified here.

## Supporting information

**S1 Fig.**
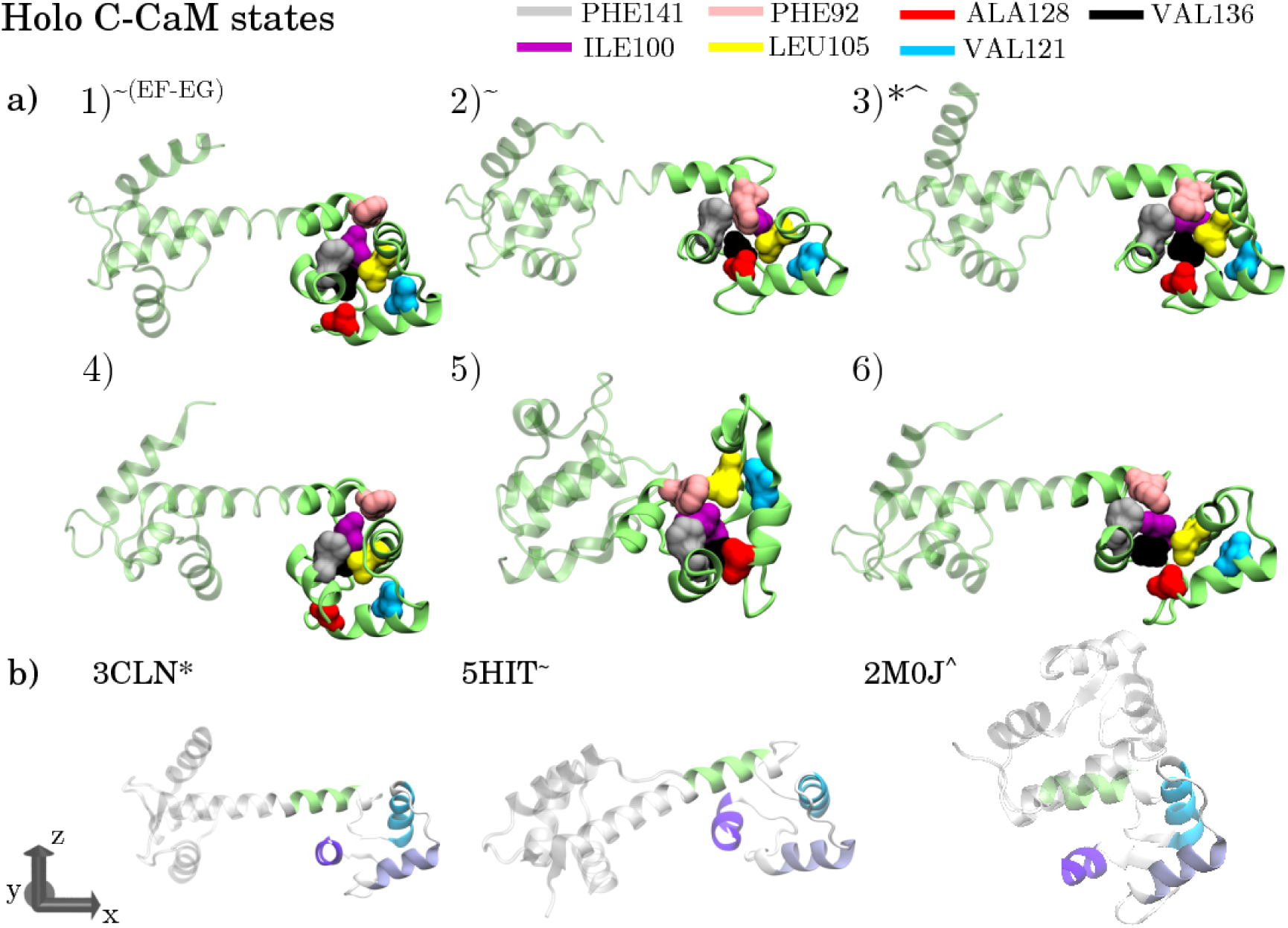
The holo C-CaM states. a) The holo C-CaM states representative structures obtained from spectral clustering. Key residues from the contact/solvent exposure analysis are highlighted in colors. b) Experimentally obtained structures with similar interhelical angle arrangements as the obtained states. The states that are similar to experimentally obtained states are marked by a symbol corresponding to the experimental structure. Note that state 1 is similar to 5HIT only at the EF-EG angles.

**S2 Fig.**
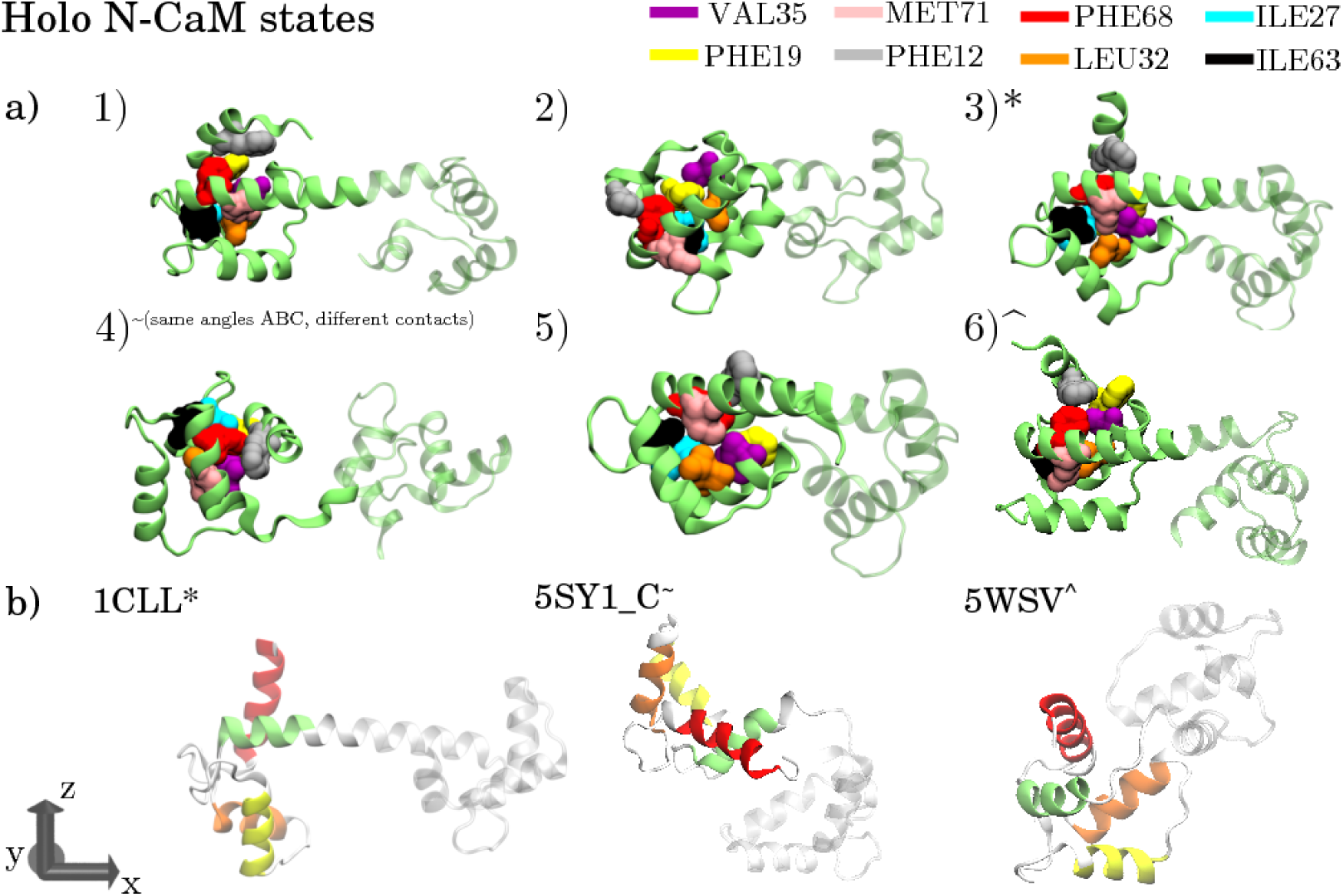
The holo N-CaM states. a) The holo N-CaM states representative structures obtained from spectral clustering. Key residues from the contact/solvent exposure analysis are highlighted in colors. b) Experimentally obtained structures with similar interhelical angle arrangements as the obtained states. The states that are similar to experimentally obtained states are marked by a symbol corresponding to the experimental structure. Note that state 4 has similar interhelical angles as 5SY1 C but a different set of inter-residue contacts.

**S3 Fig.**
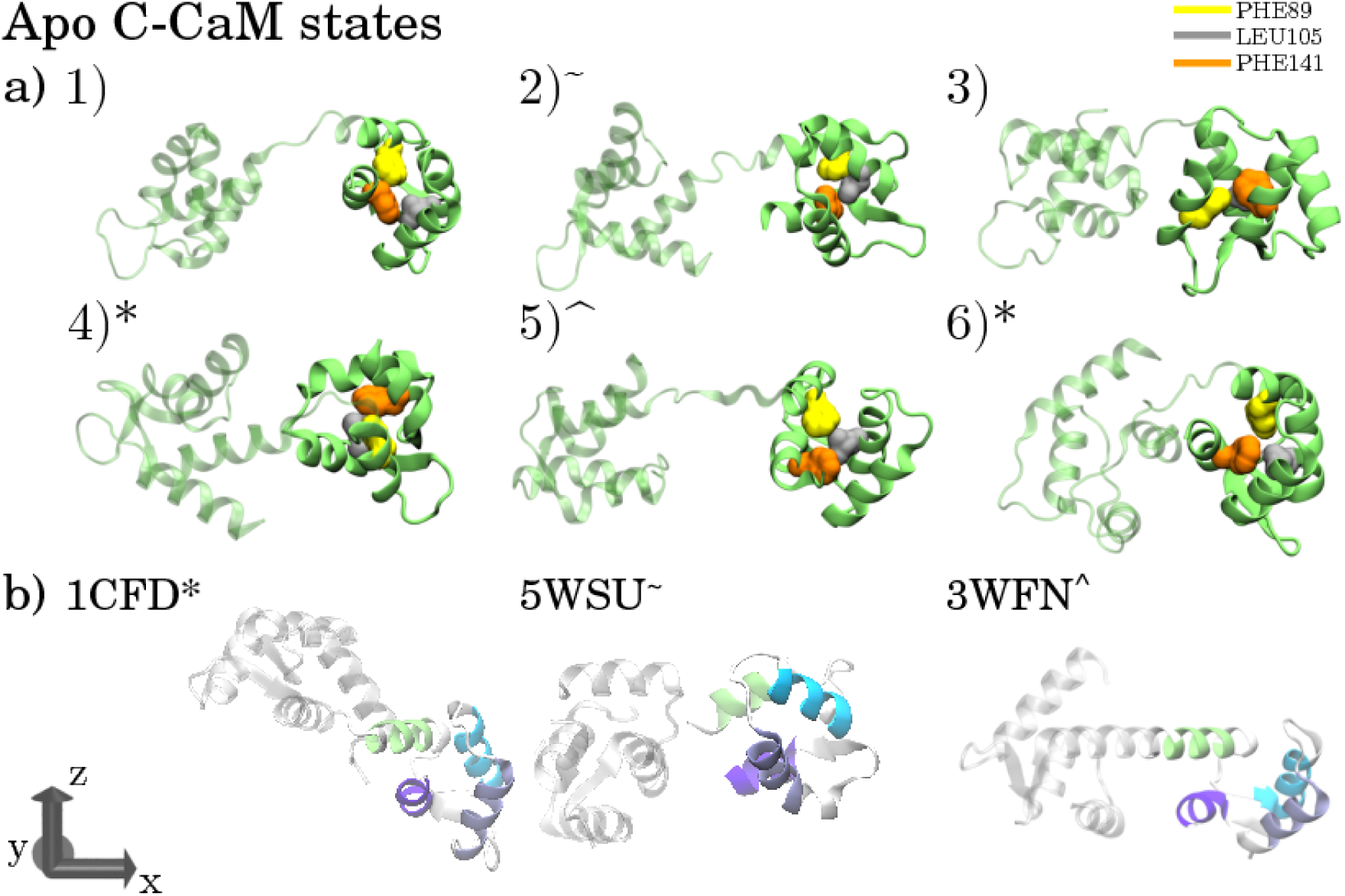
The apo C-CaM states. a) The apo C-CaM states representative structures obtained from spectral clustering. Key residues from the contact/solvent exposure analysis are highlighted in colors. b) Experimentally obtained structures with similar interhelical angle arrangements as the obtained states. The states that are similar to experimentally obtained states are marked by a symbol corresponding to the experimental structure.

**S4 Fig.**
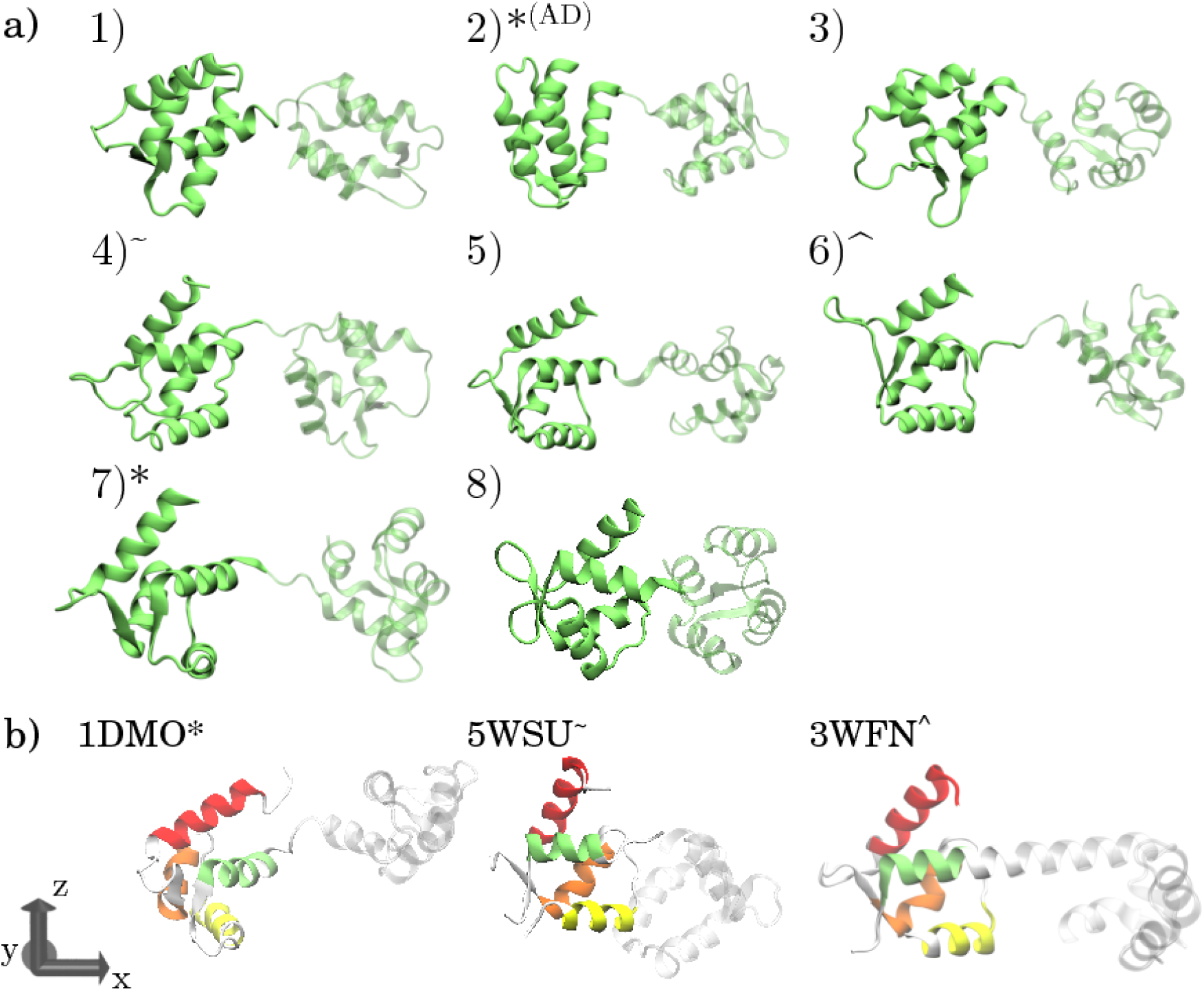
The apo N-CaM states. a) The apo N-CaM states representative structures obtained from spectral clustering. b) Experimentally obtained structures with similar interhelical angle arrangements as the obtained states. The states that are similar to experimentally obtained states are marked by a symbol corresponding to the experimental structure. Note that state 2 is similar to 1DMO only for the AD angle.

**S5 Fig.**
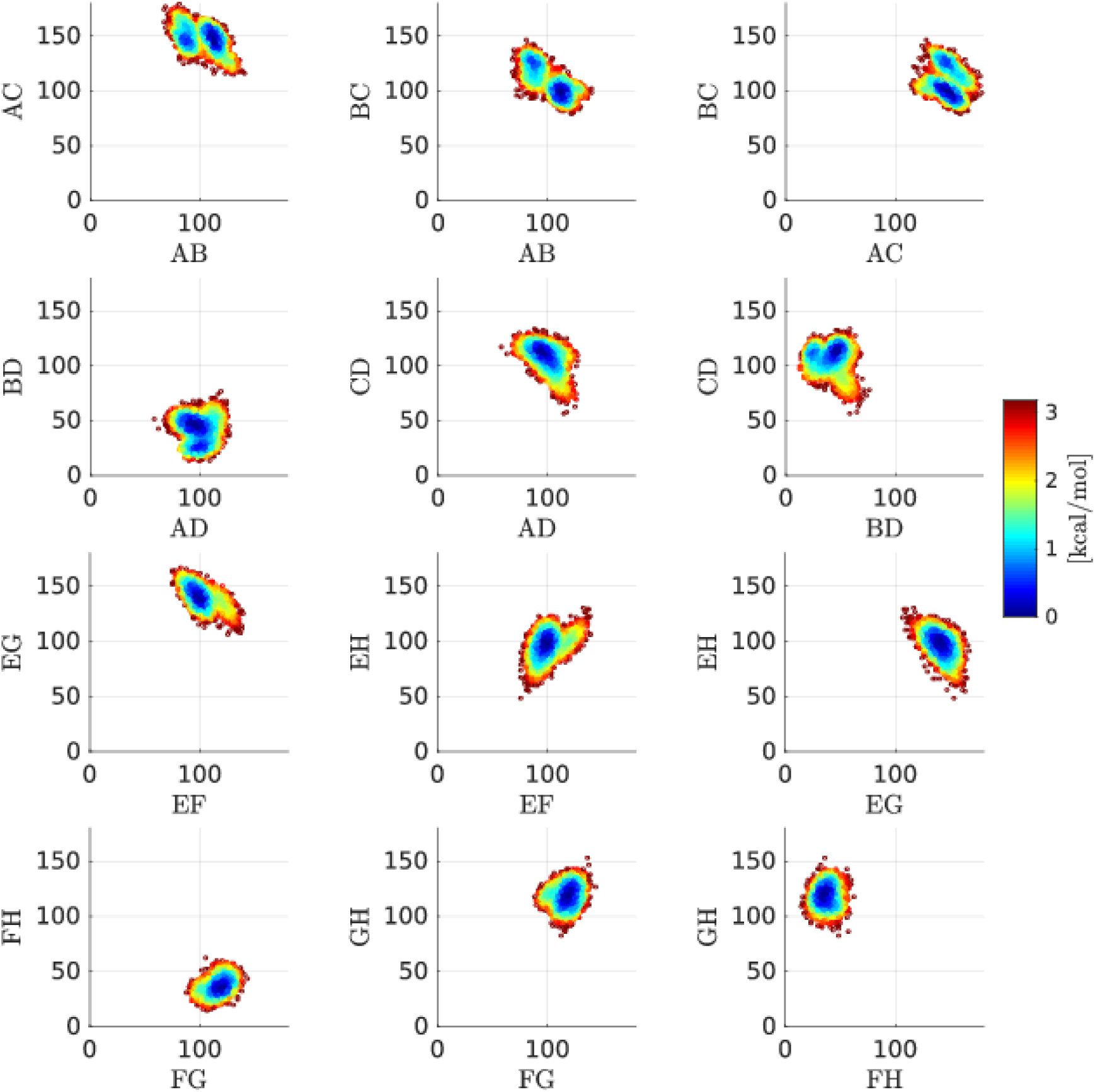
Estimated free energy landscapes of the holo ensemble using data acquired from MD simulations. Note that these are not accurate estimates of free energies due to limited simulation time.

**S6 Fig.**
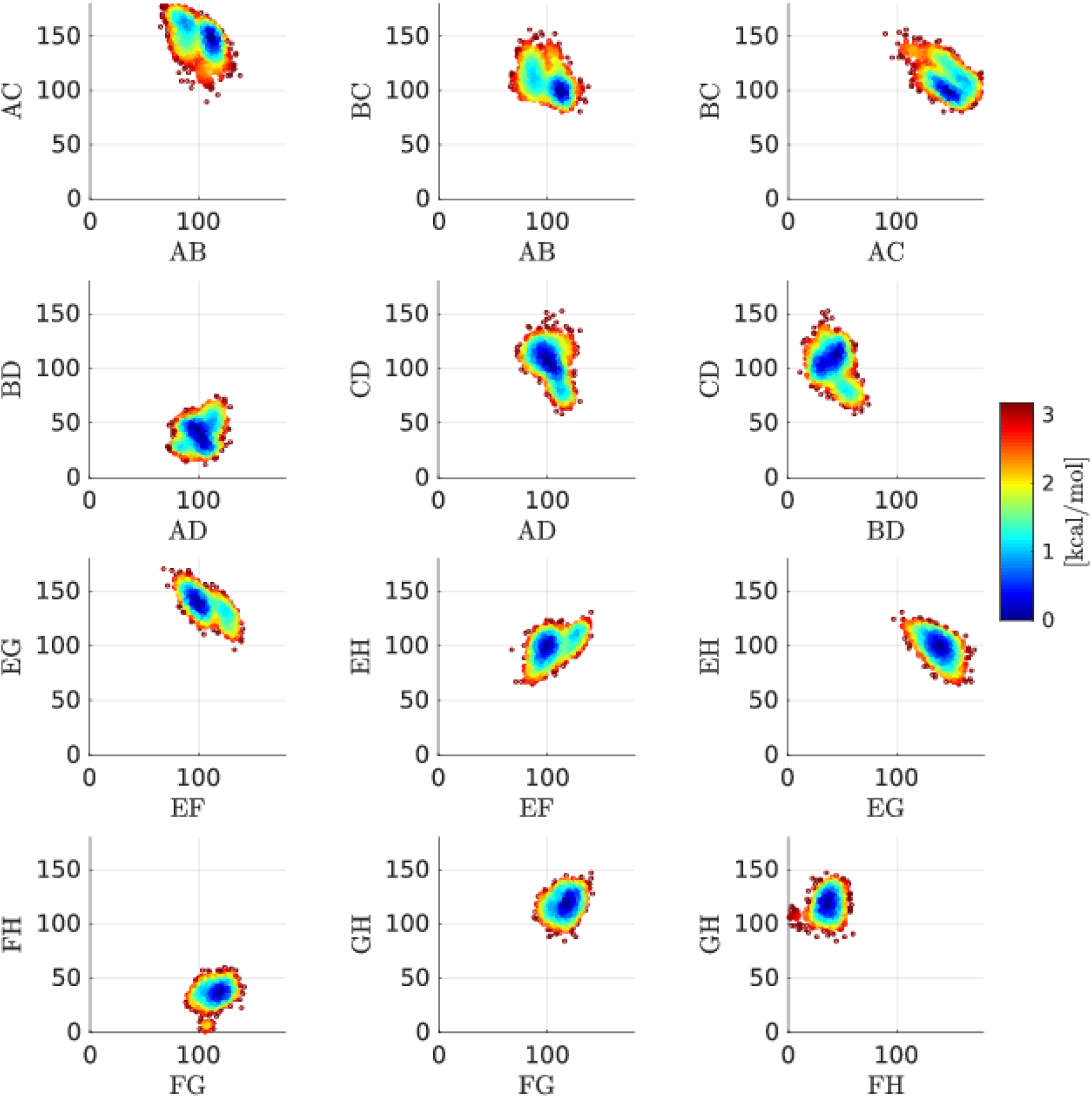
Estimated free energy landscapes of the holo ensemble using data acquired from T-REMD simulations. Note that these are not accurate estimates of free energies due to limited simulation time.

**S7 Fig.**
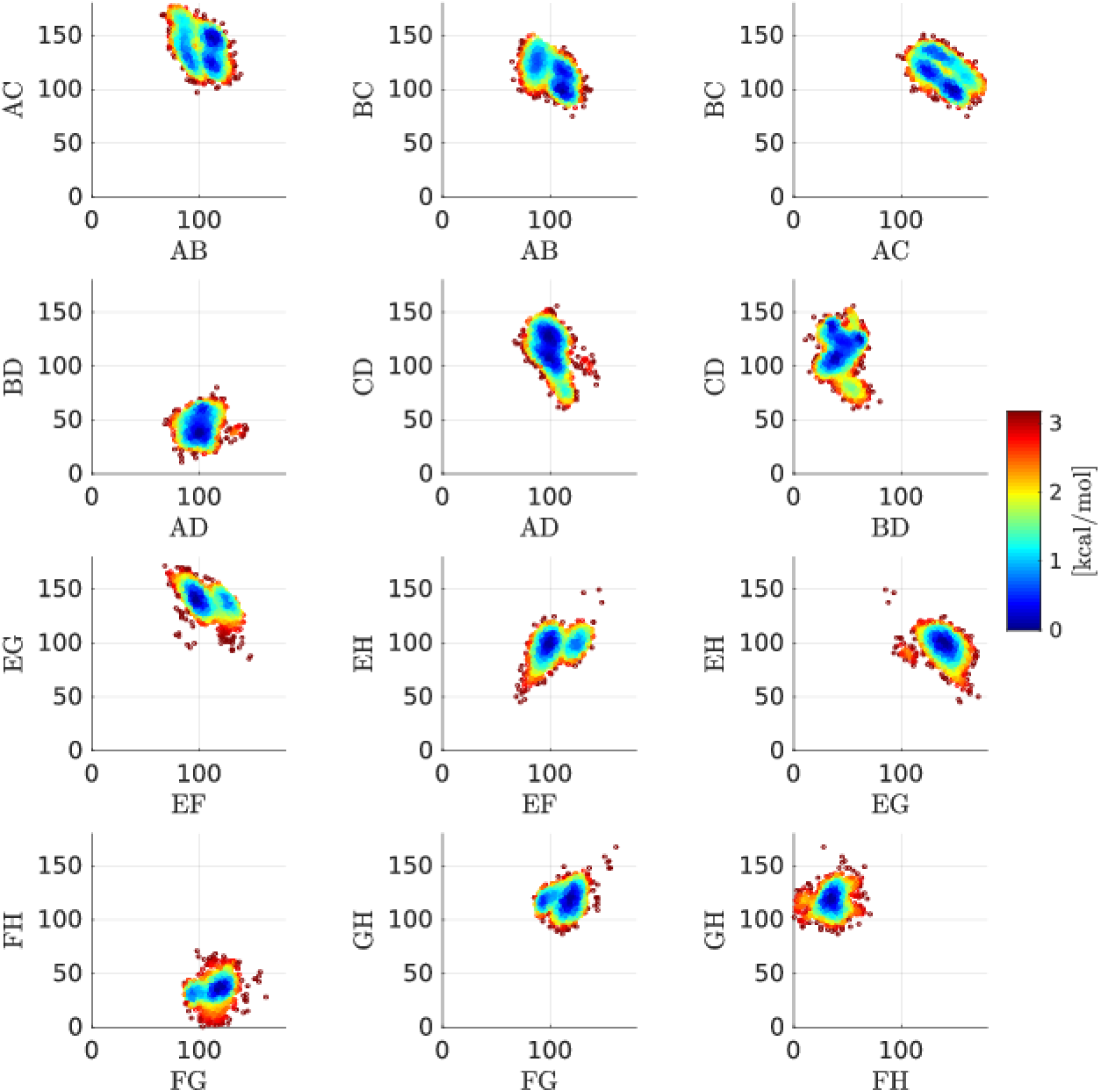
Estimated free energy landscapes of the holo ensemble using data acquired from REST simulations. Note that these are not accurate estimates of free energies due to limited simulation time.

**S8 Fig.**
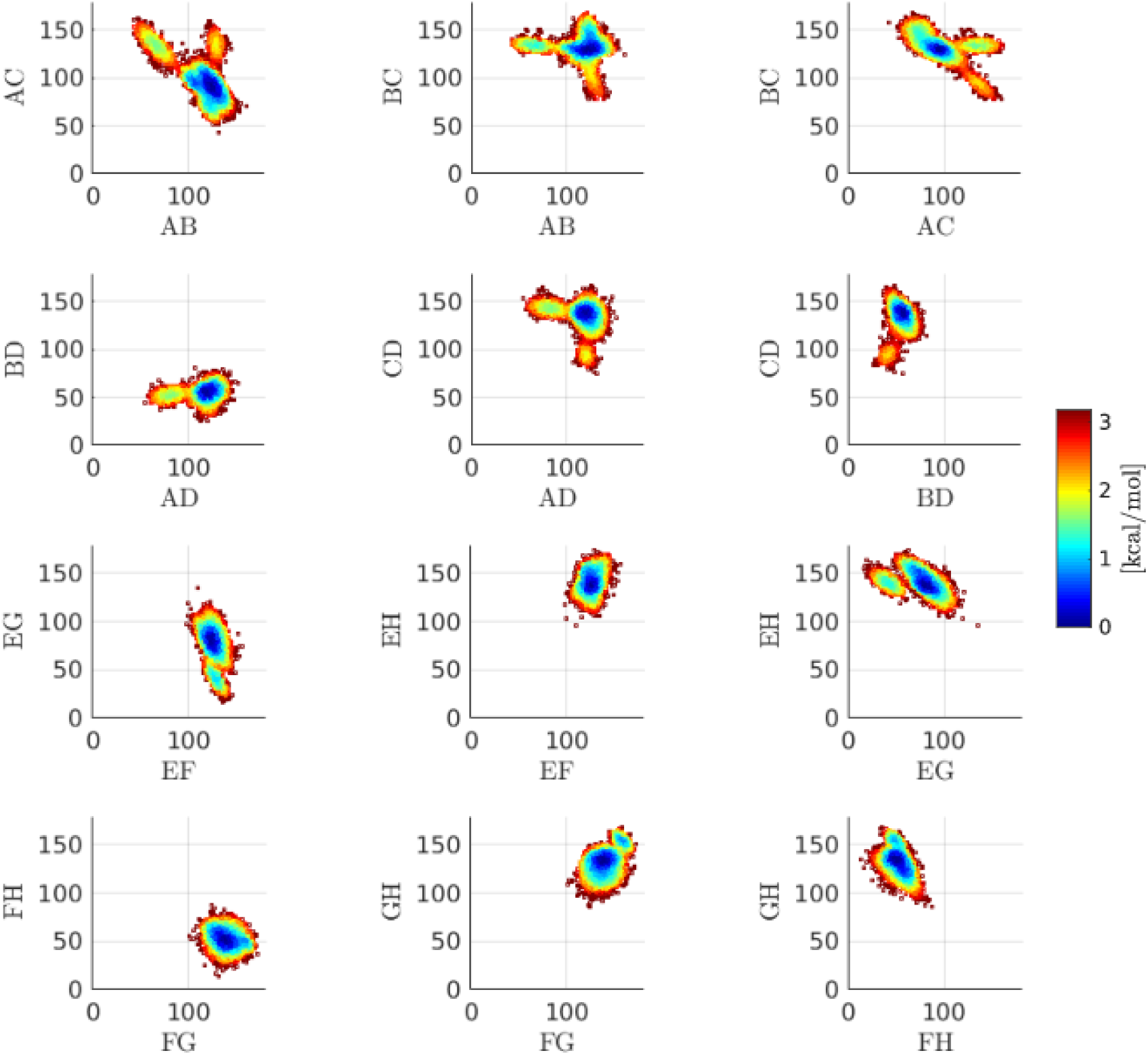
Estimated free energy landscapes of the apo ensemble using data acquired from MD simulations. Note that these are not accurate estimates of free energies due to limited simulation time.

**S9 Fig.**
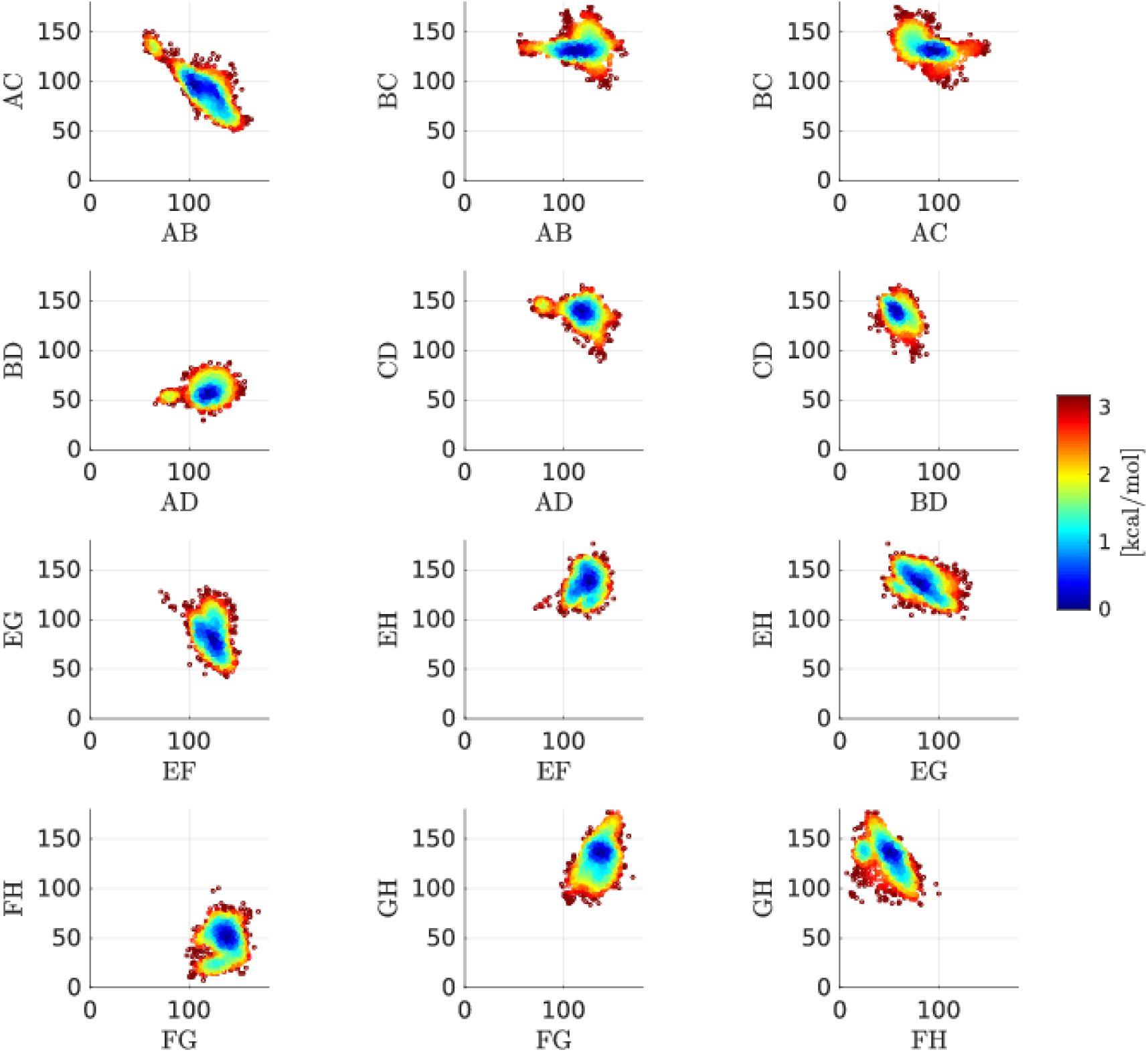
Estimated free energy landscapes of the apo ensemble using data acquired from T-REMD simulations. Note that these are not accurate estimates of free energies due to limited simulation time.

**S10 Fig.**
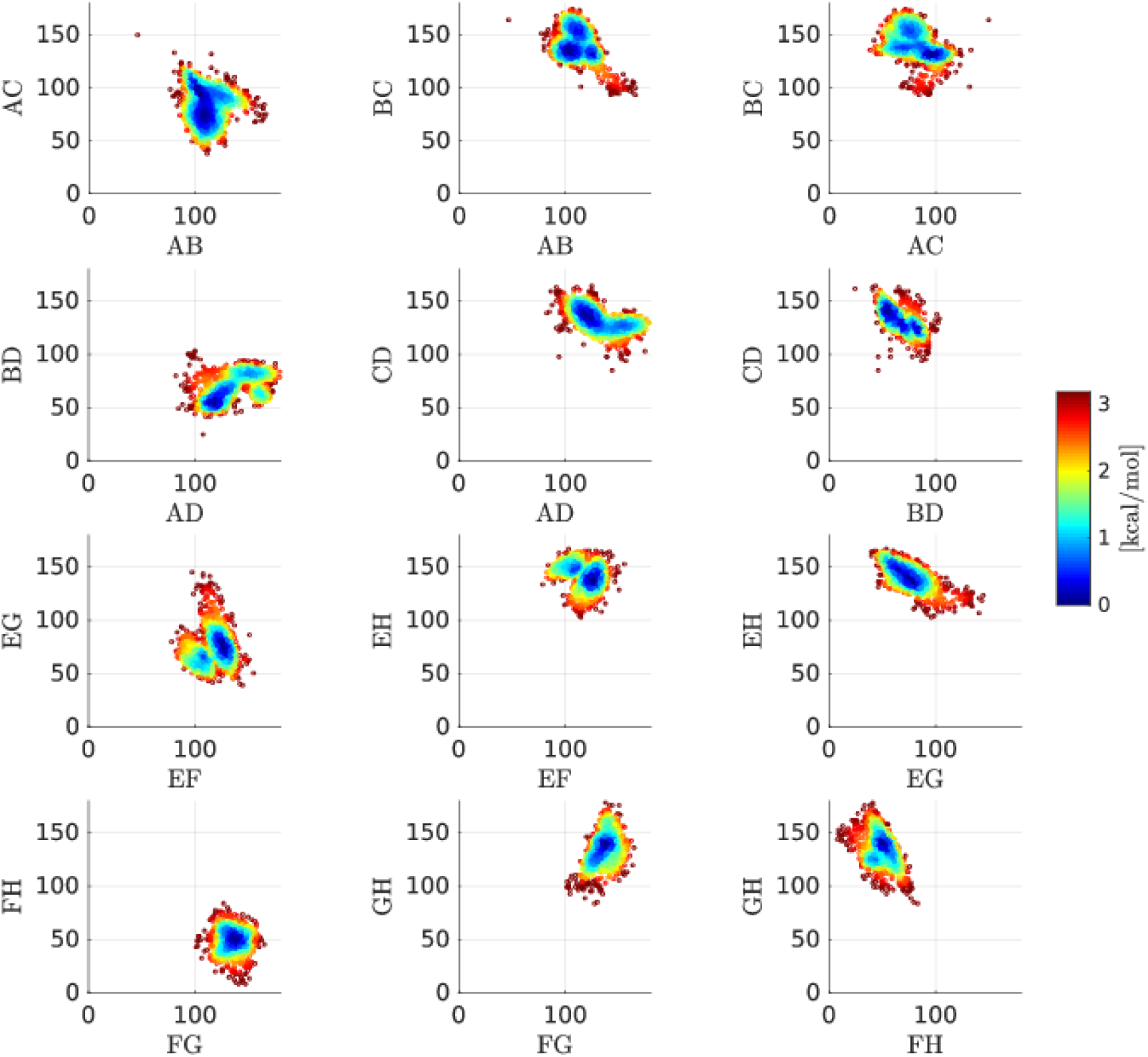
Estimated free energy landscapes of the apo ensemble using data acquired from REST simulations. Note that these are not accurate estimates of free energies due to limited simulation time.

**S11 Fig.**
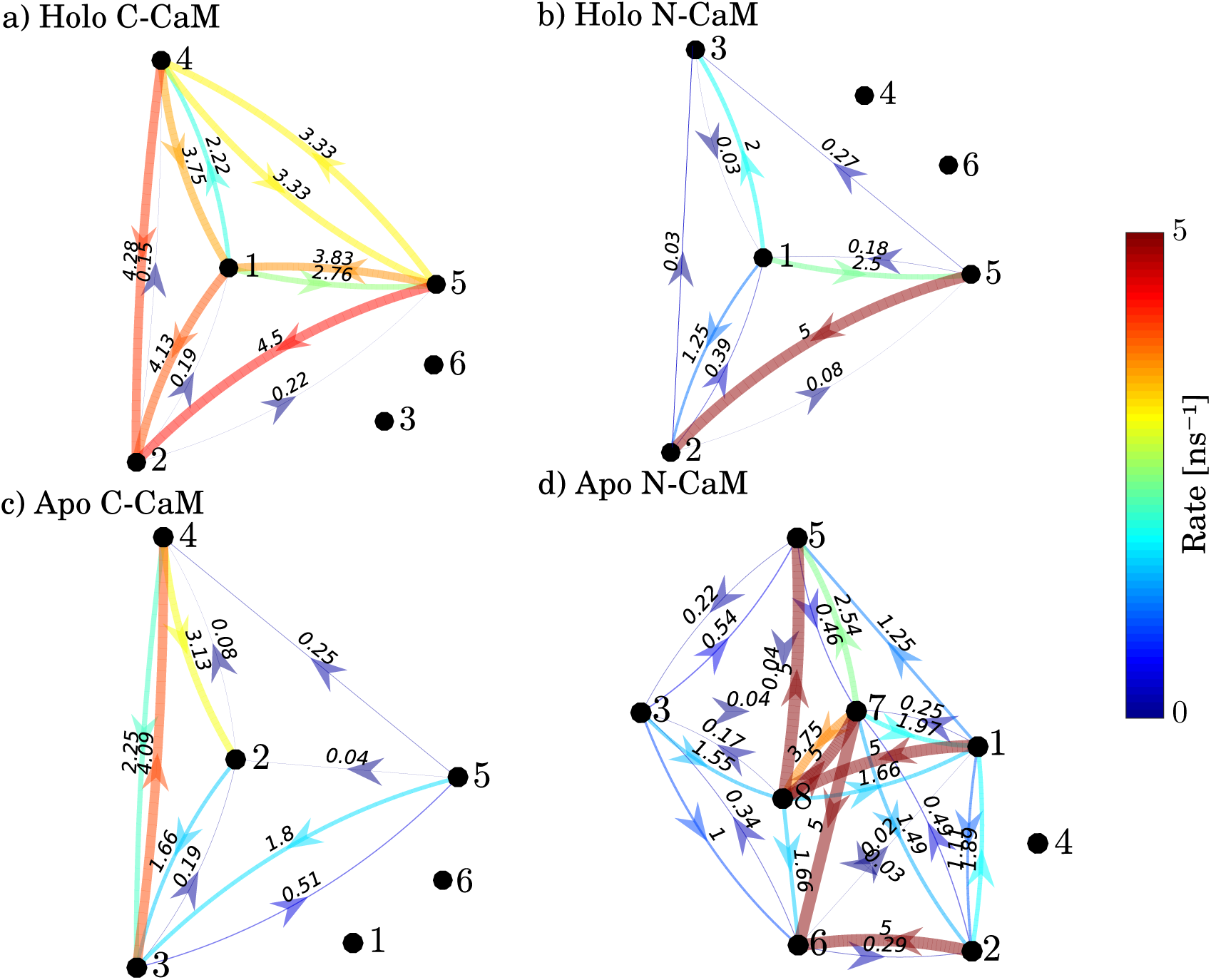
Networks of interconversion between the states obtained in this study, as derived from the MD simulation trajectories. States that were not sampled in the plain MD simulations are disconnected from the networks. Replica exchange simulations disrupt the dynamics through coordinate exchanges. Although kinetics of toy systems can be restored from replica exchange simulations [90], application of the method to the present dataset was not possible.

**S12 Fig.**
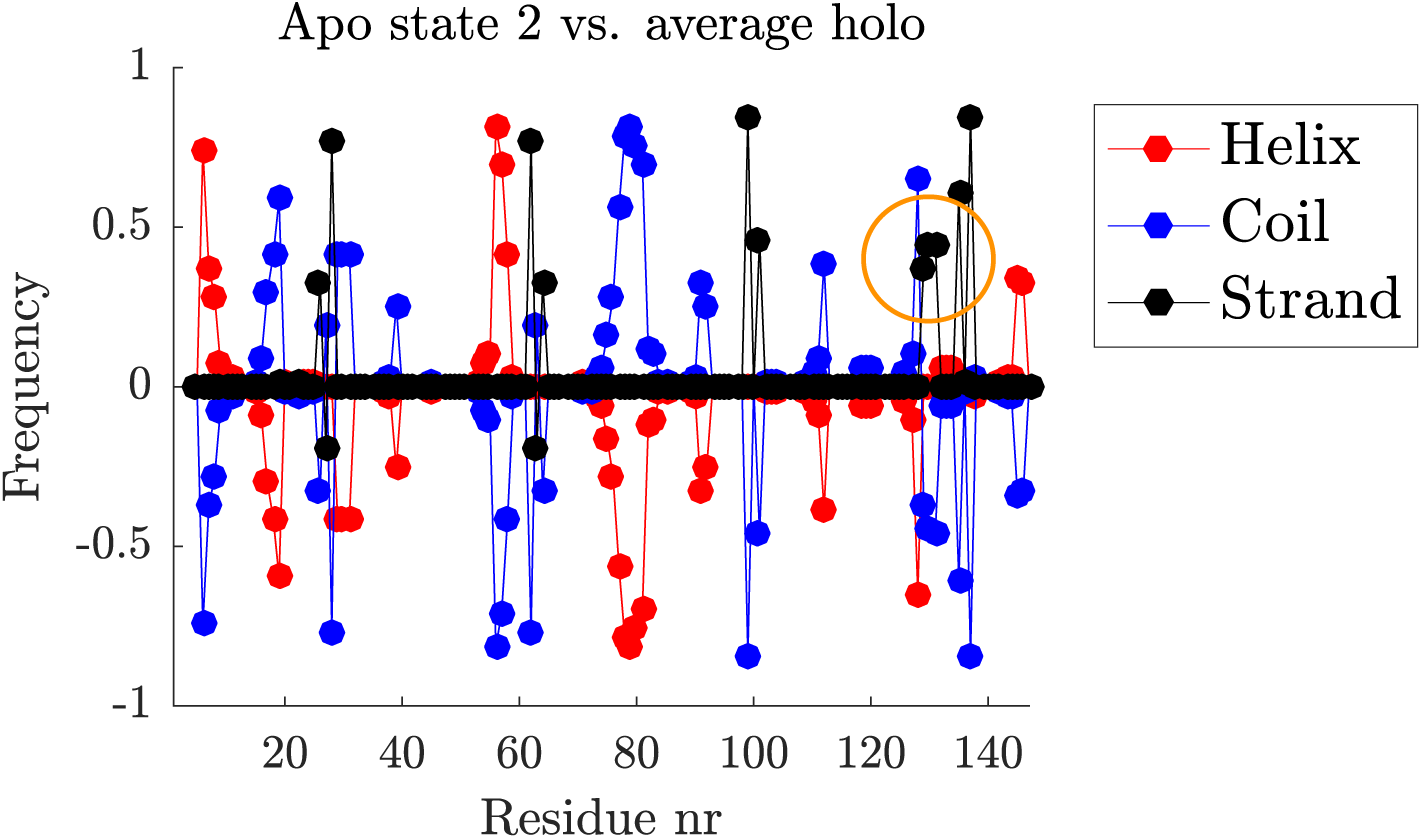
Secondary structure frequency of apo state 2 difference to average holo secondary structure frequency. The propensity of residues 129-131 to join the beta sheet and deforming the fourth Ca^2+^-loop is marked by an orange circle.

**S13 Fig.**
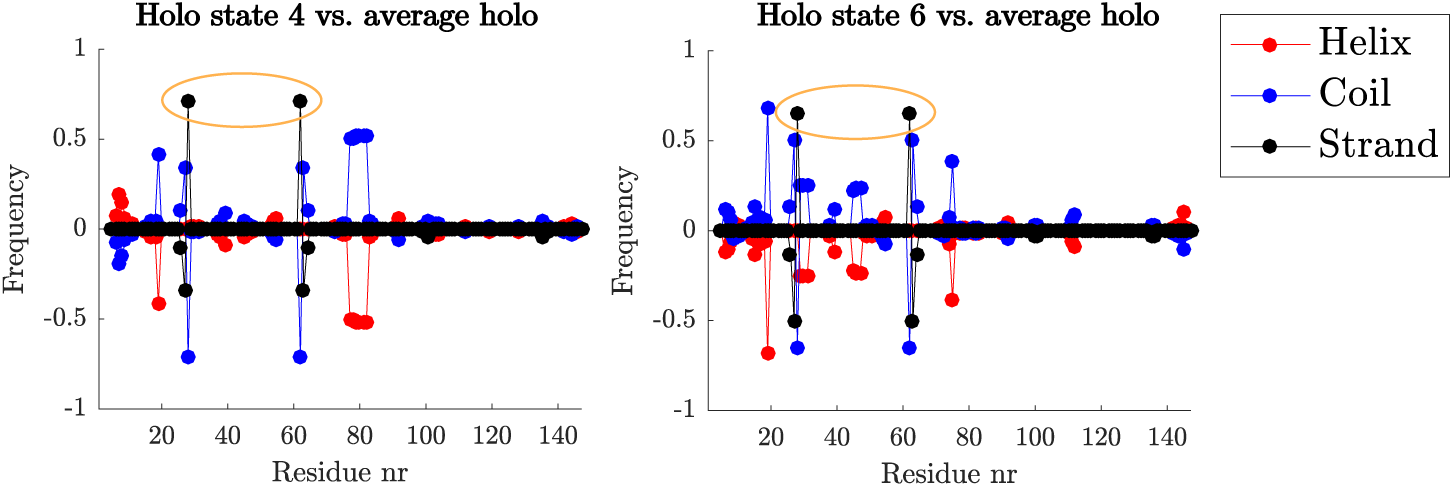
Secondary structure frequency of holo N-CaM state 4 and 6 difference to average holo secondary structure frequency. These states overlap the apo ensemble. The beta sheet shift to residue 28/62 in N-CaM is marked by an orange ellipse.

**S14 Fig.**
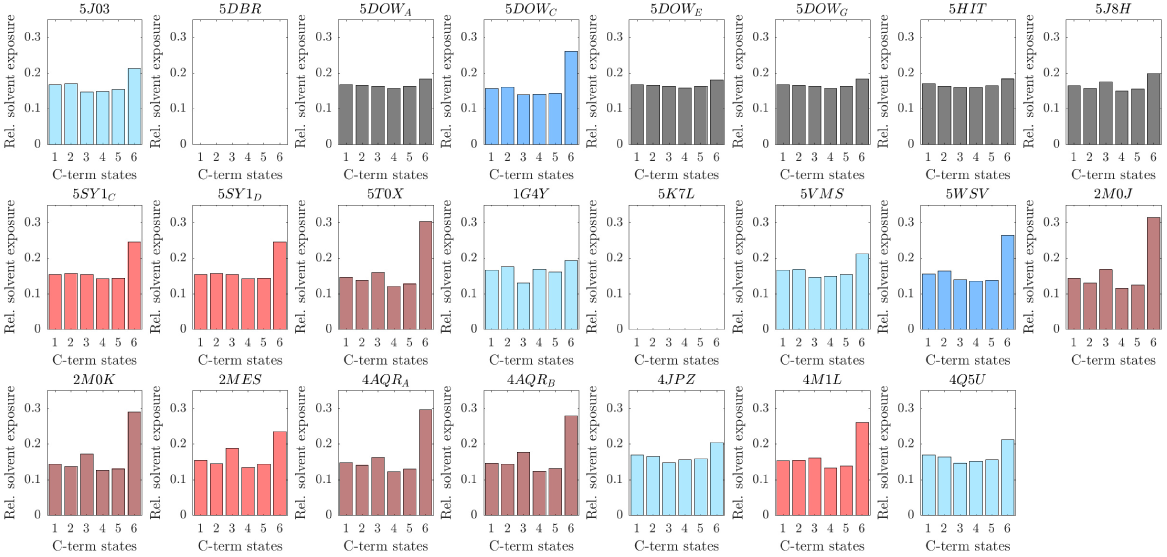
Holo C-CaM total relative solvent exposure for each CaM-complex structure.

**S15 Fig.**
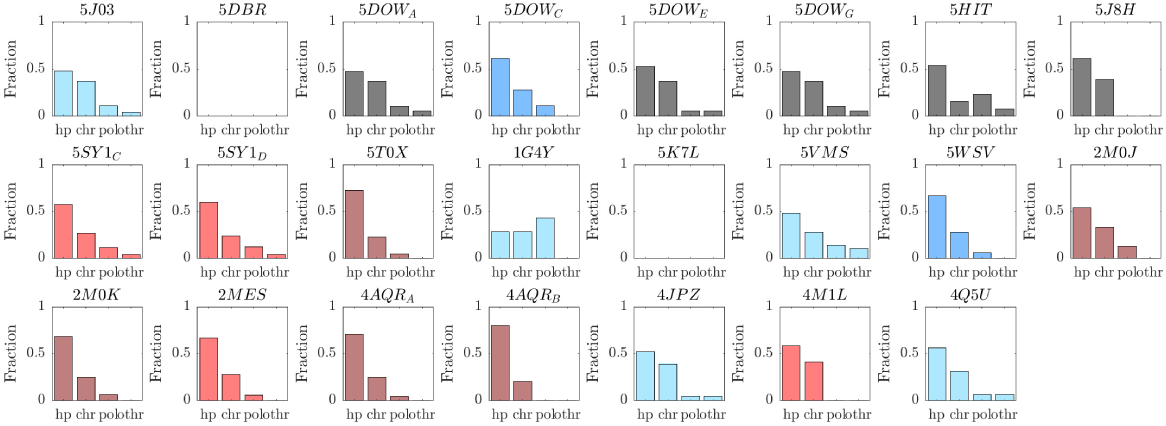
The residue types involved in holo C-CaM target protein contacts for each CaM-complex structure.

**S16 Fig.**
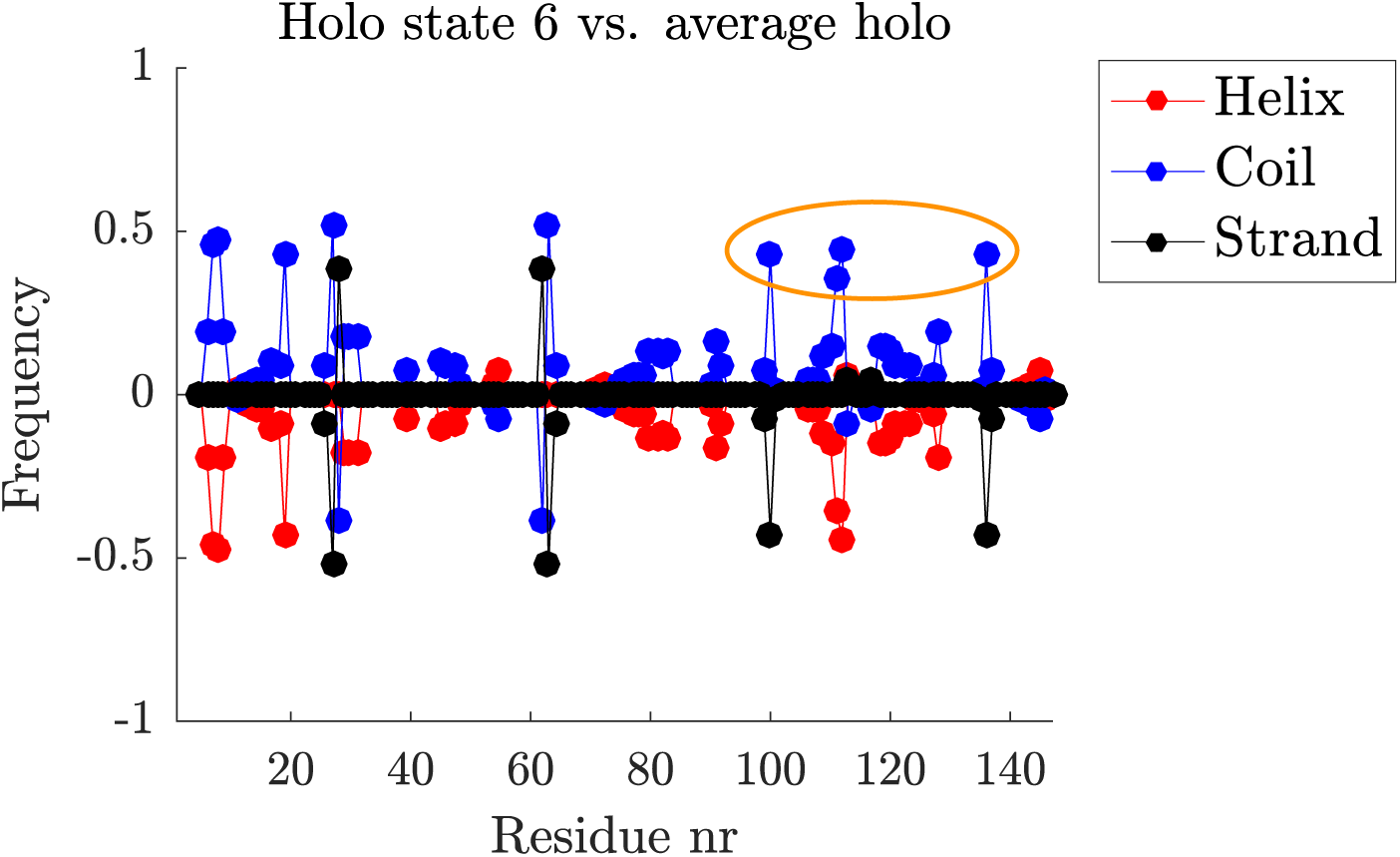
Secondary structure frequency of holo state 6 difference to average holo secondary structure frequency. The lack of beta sheets in the C-term lobe is marked by an orange ellipse.

**S17 Fig.**
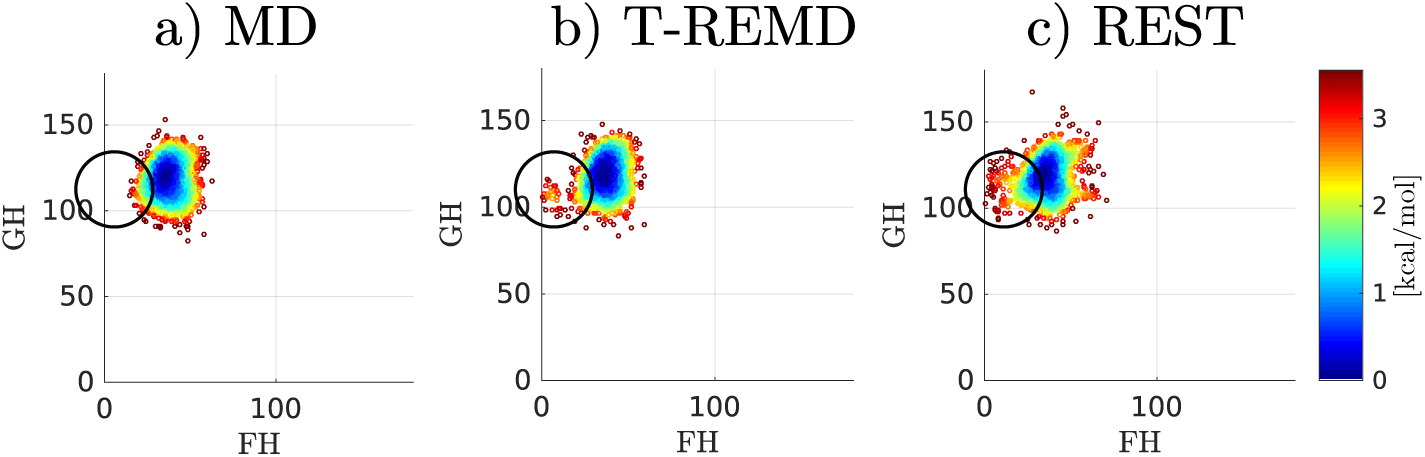
Free energy landscapes over holo FH and GH interhelical angles, estimated with GMM with cross validation free energy estimator [43]. State 6 is marked with a ring, showing that it is only observed in temperature enhanced MD.

**S18 Fig.**
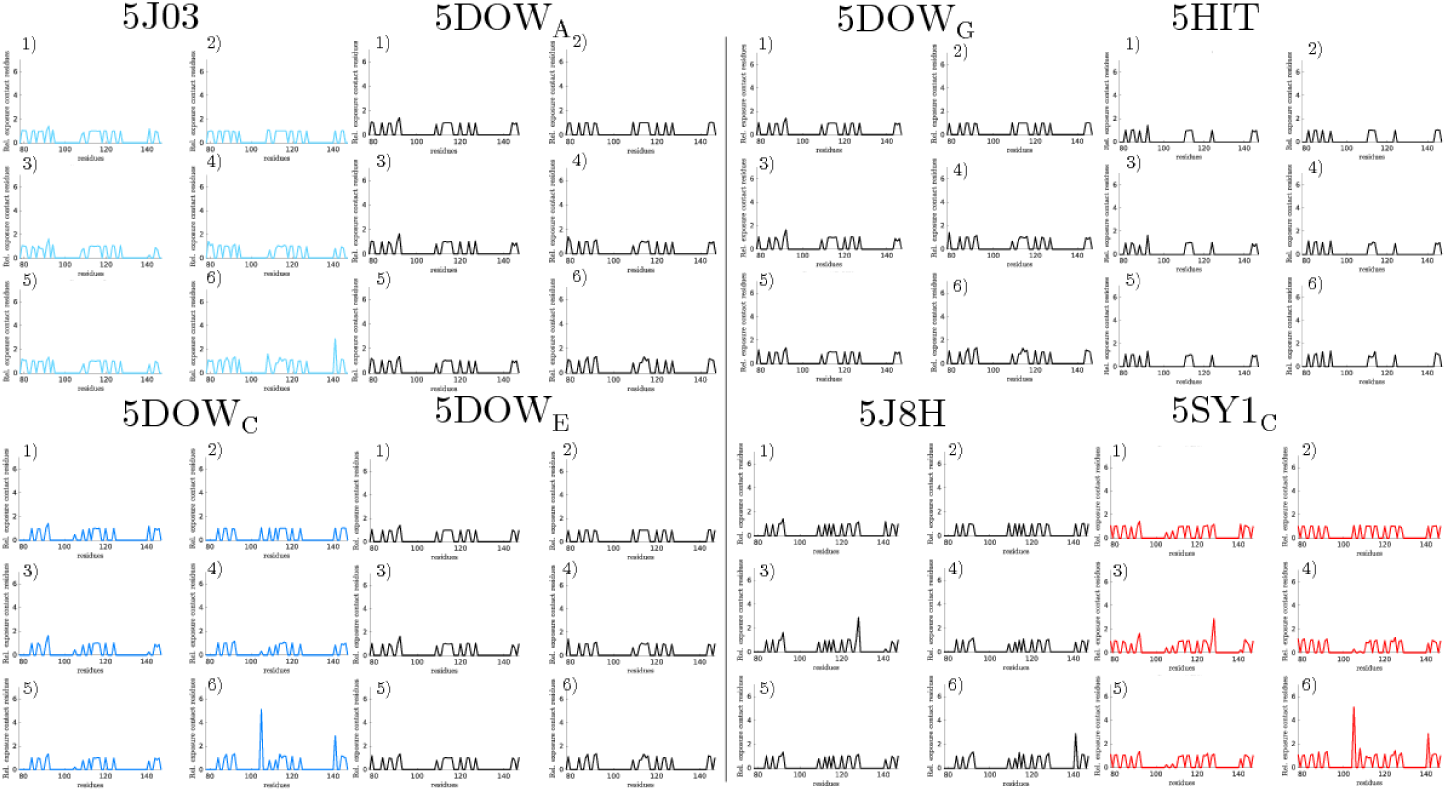
The relative solvent exposure per CaM-complex contact residue for the different holo C-CaM states, part 1.

**S19 Fig.**
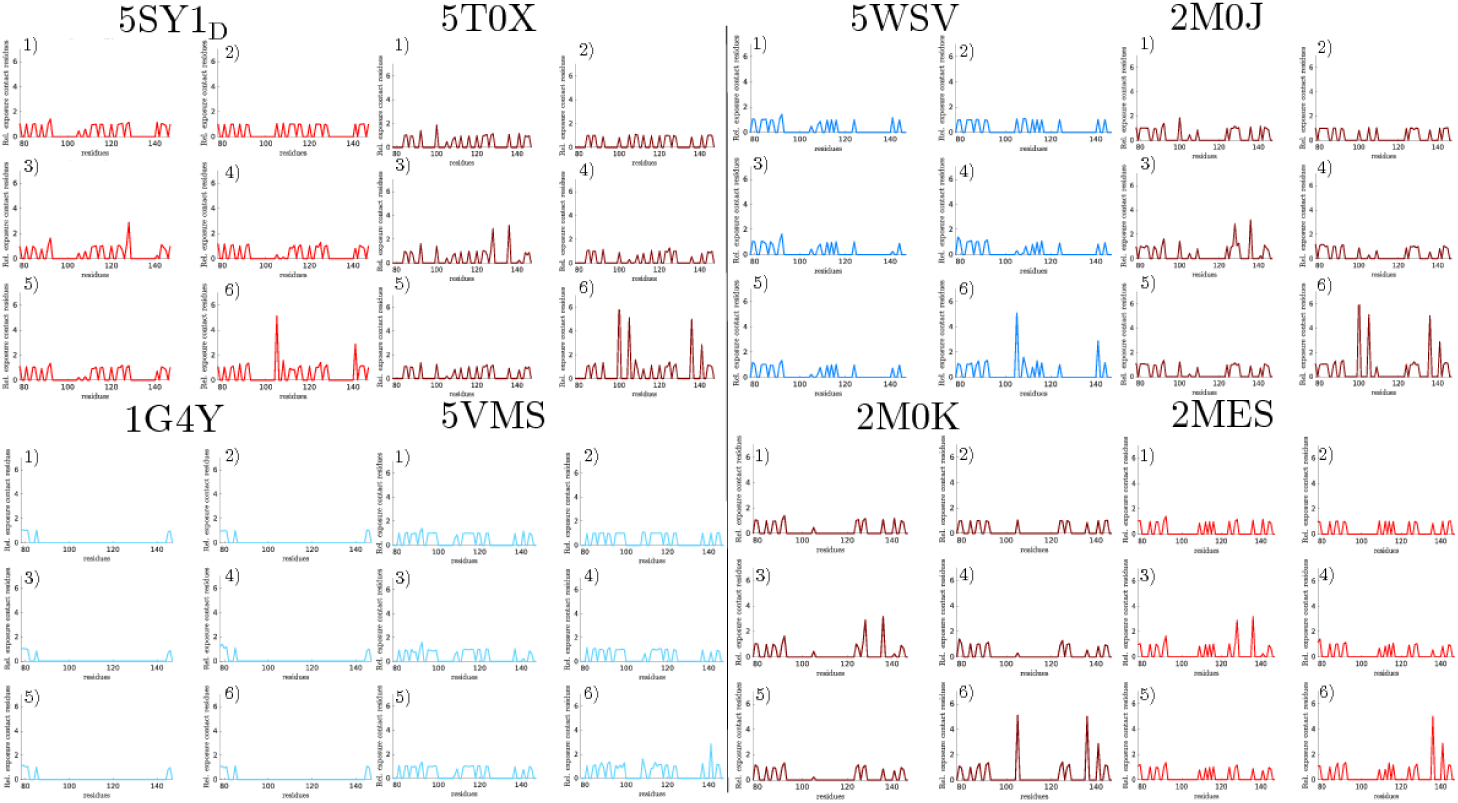
The relative solvent exposure per CaM-complex contact residue for the different holo C-CaM states, part 2.

**S20 Fig.**
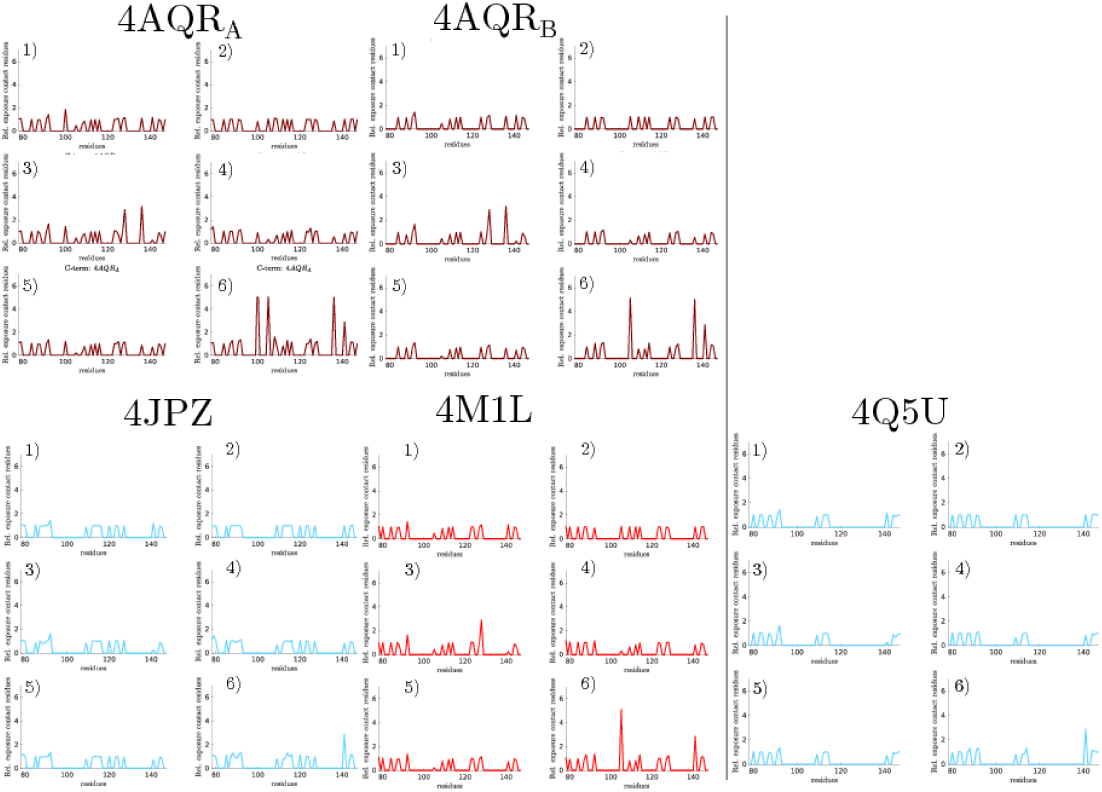
The relative solvent exposure per CaM-complex contact residue for the different holo C-CaM states, part 3.

**S21 Fig.**
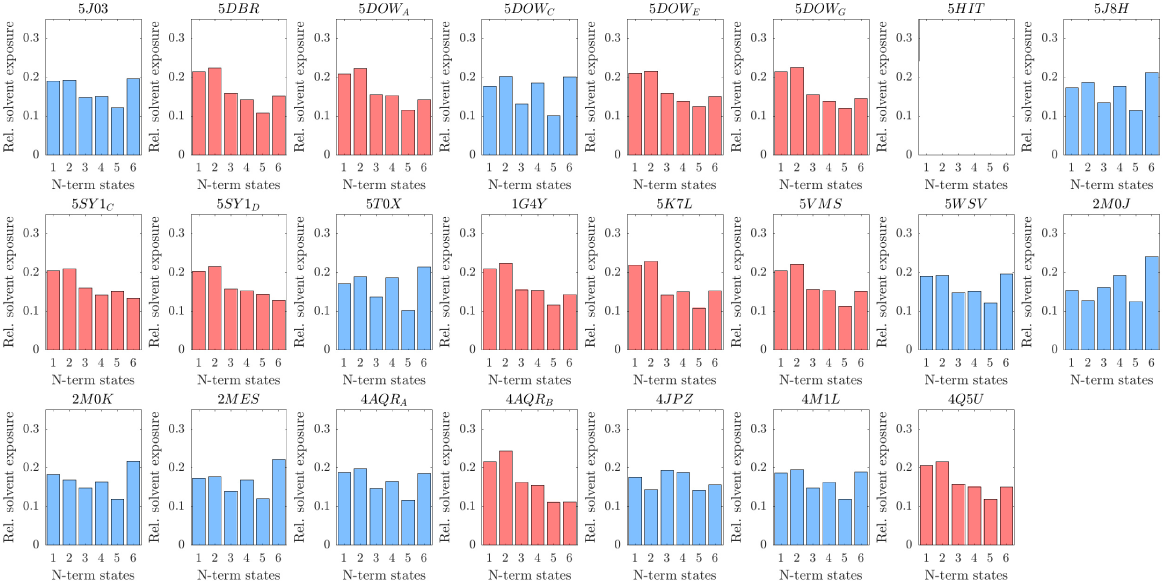
Holo N-CaM total relative solvent exposure for each CaM-complex structure.

**S22 Fig.**
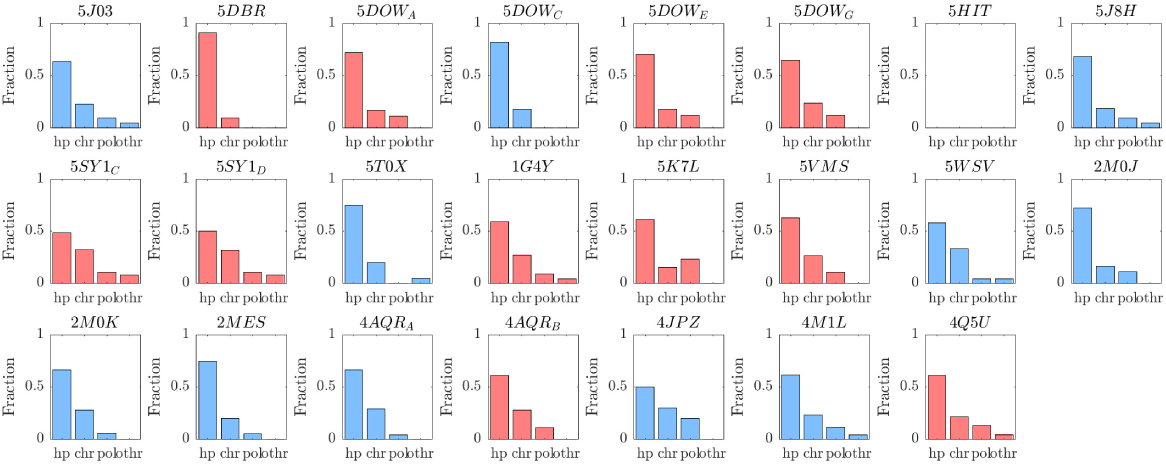
The residue types involved in holo N-CaM target protein contacts for each CaM-complex structure.

**S23 Fig.**
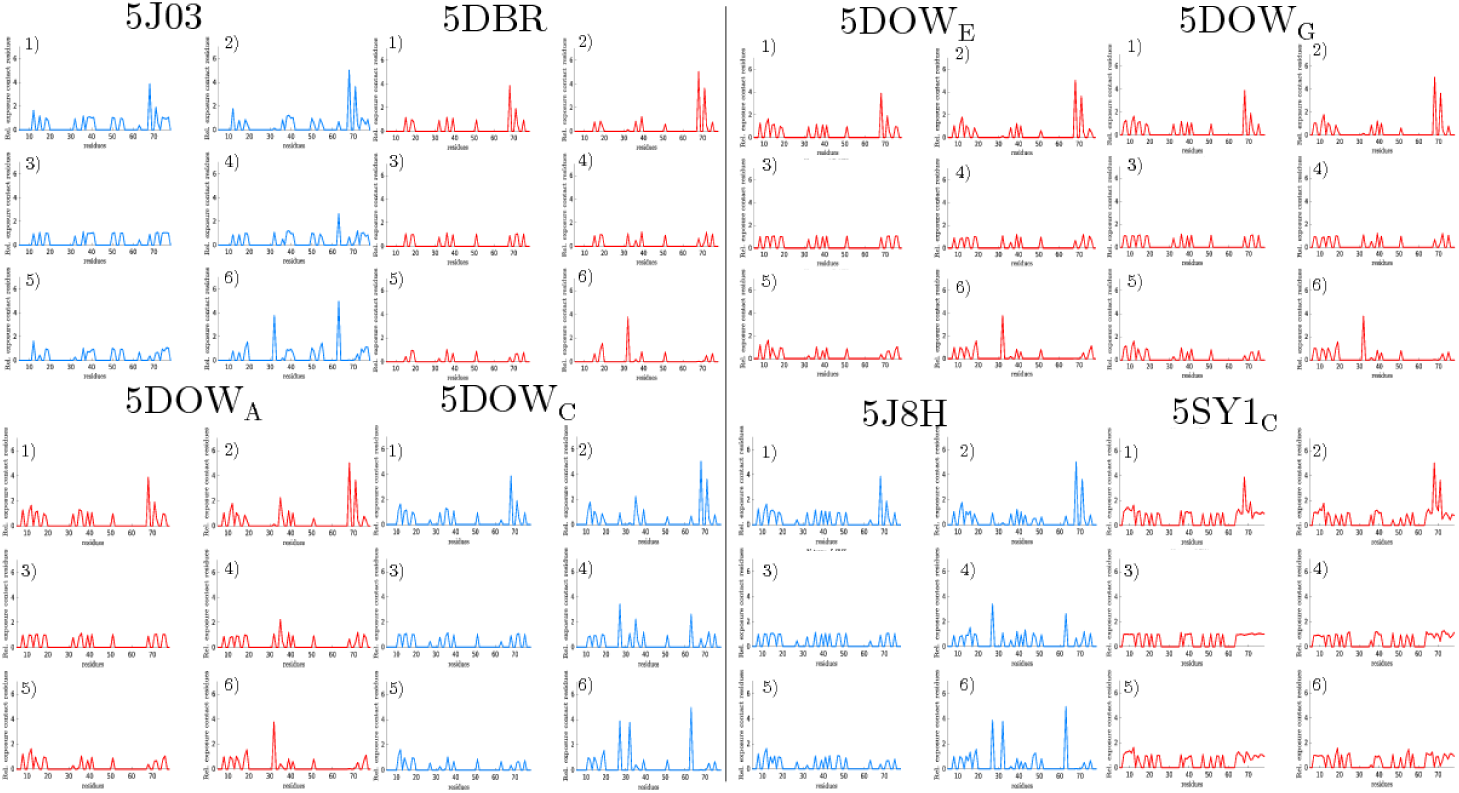
The relative solvent exposure per CaM-complex contact residue for the different holo N-CaM states, part 1.

**S24 Fig.**
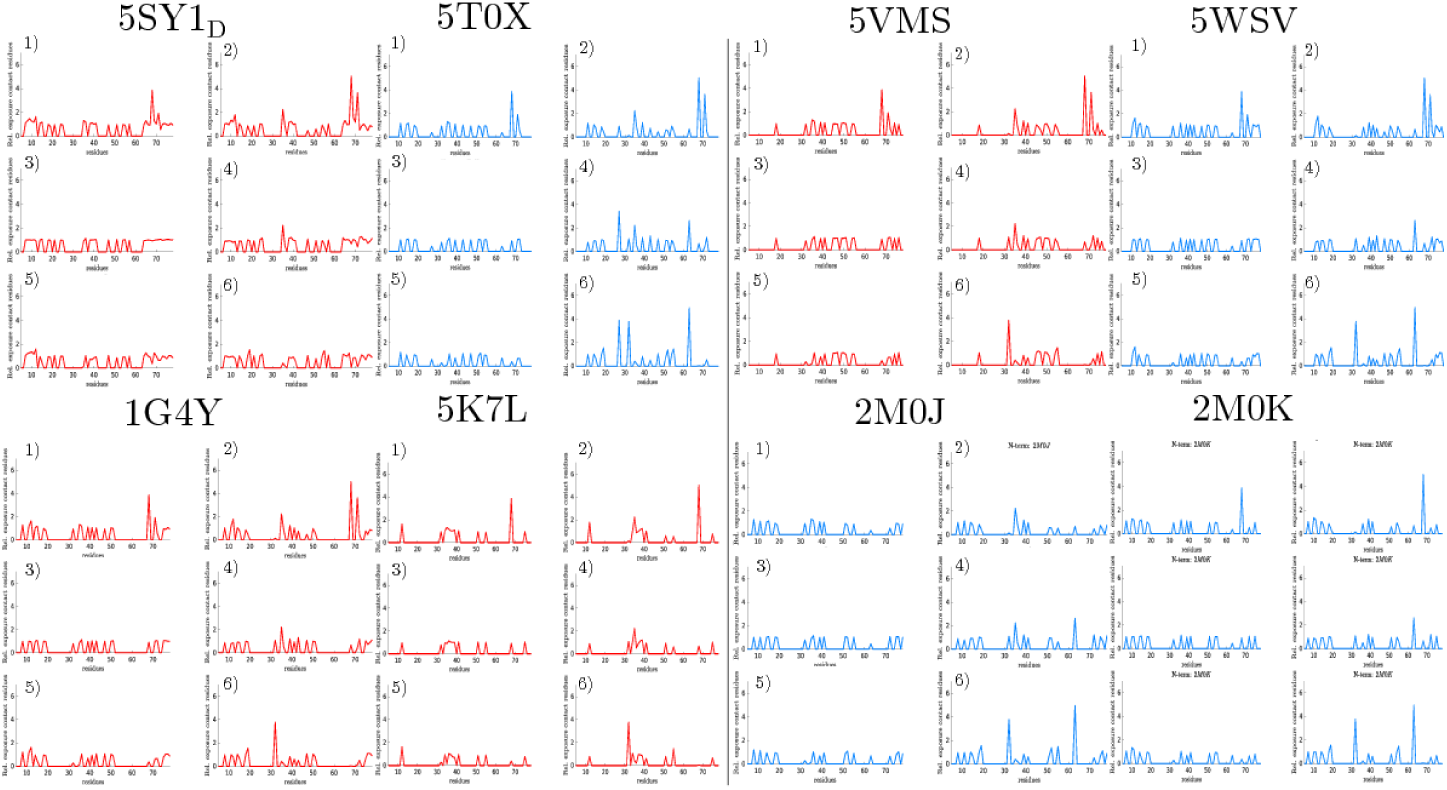
The relative solvent exposure per CaM-complex contact residue for the different holo N-CaM states, part 2.

**S25 Fig.**
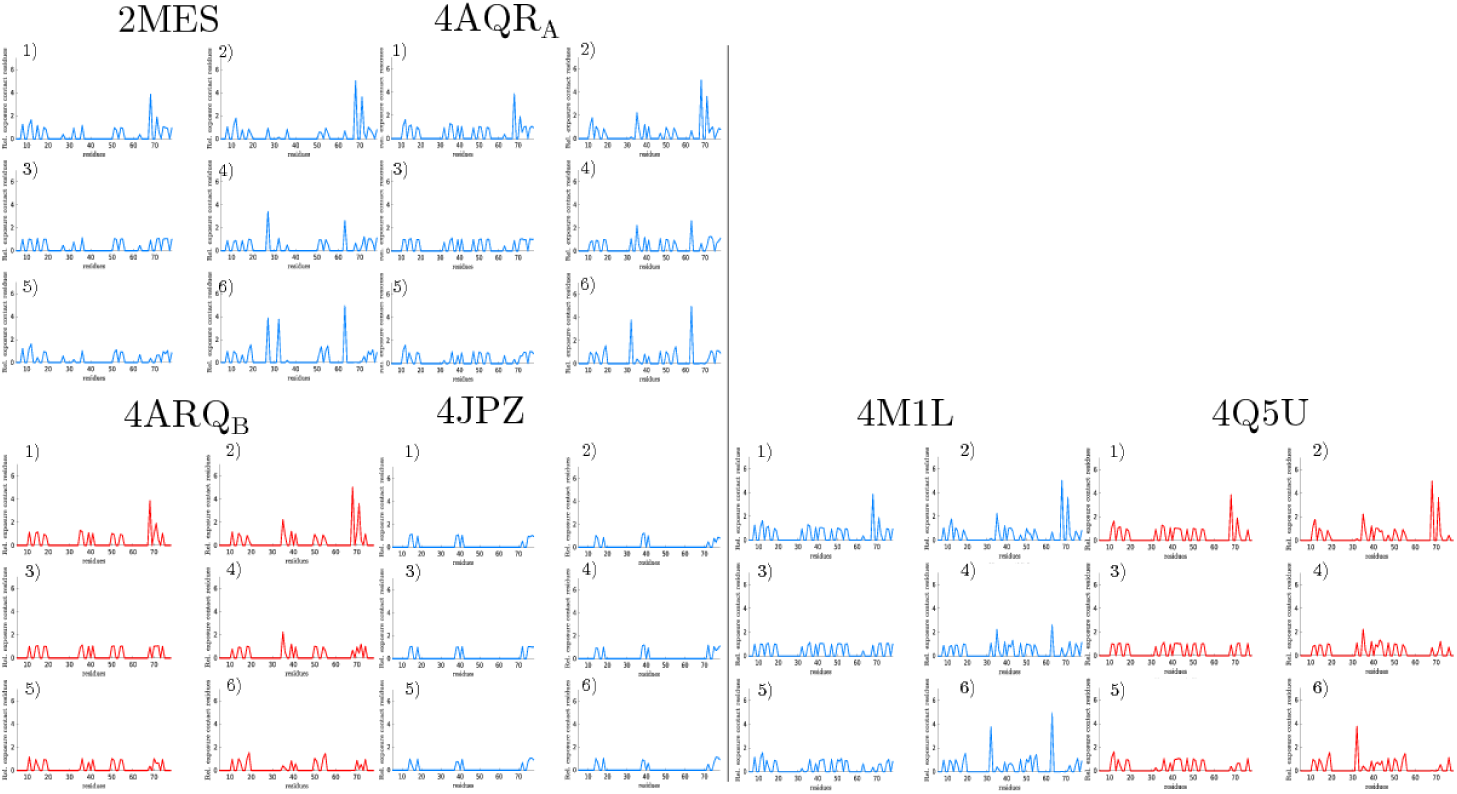
The relative solvent exposure per CaM-complex contact residue for the different holo N-CaM states, part 3.

**S26 Fig.**
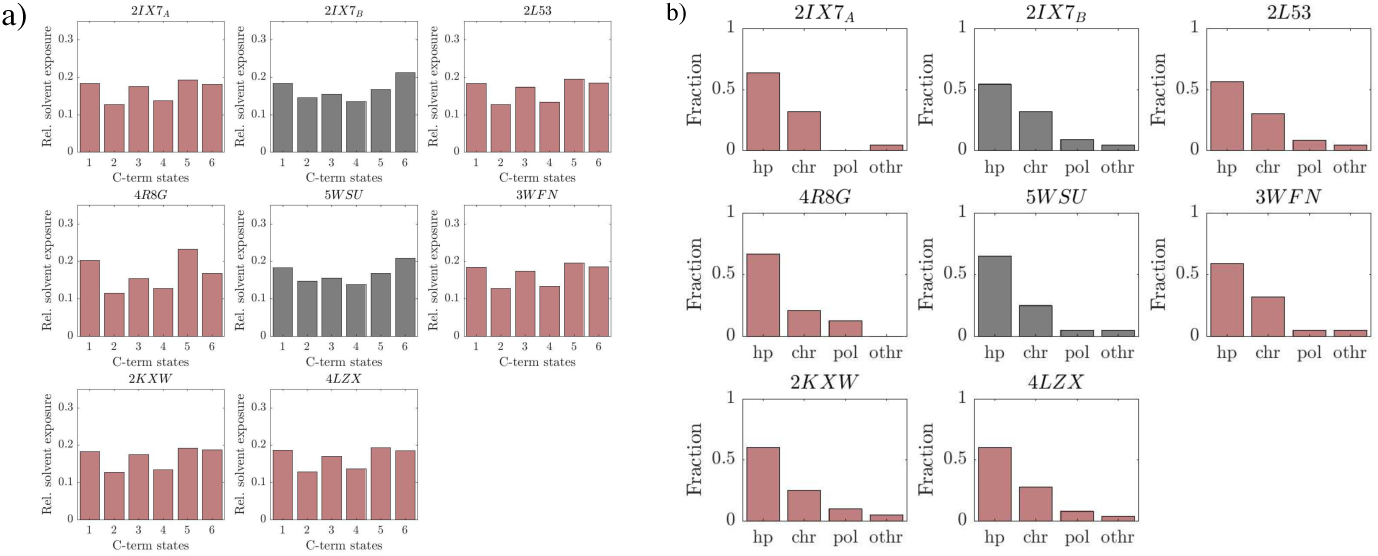
a) Apo C-CaM total relative solvent exposure and b) the residue types involved in apo C-CaM target protein contacts for each CaM-complex structure.

**S27 Fig.**
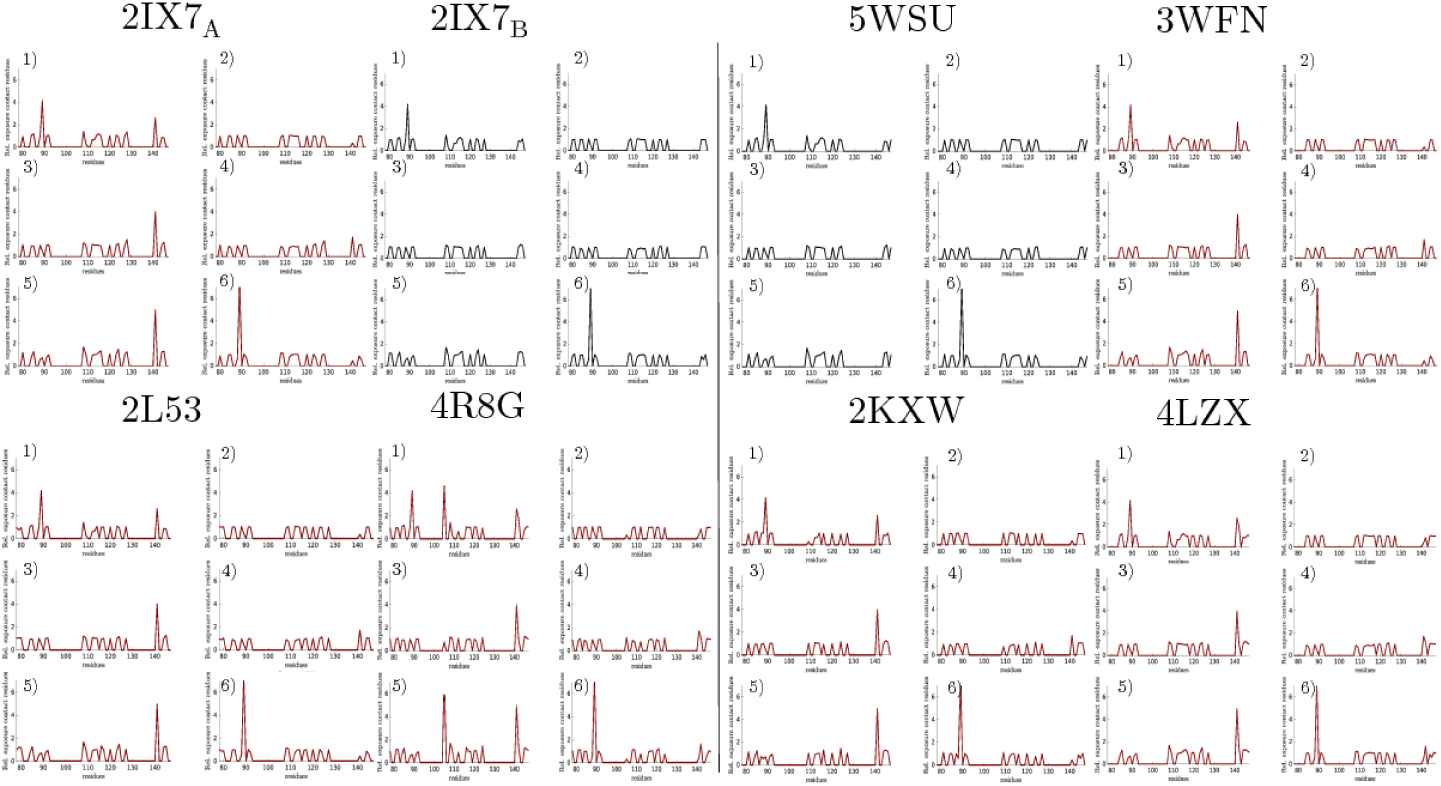
The relative solvent exposure per CaM-complex contact residue for the different apo C-CaM states.

**S28 Fig.**
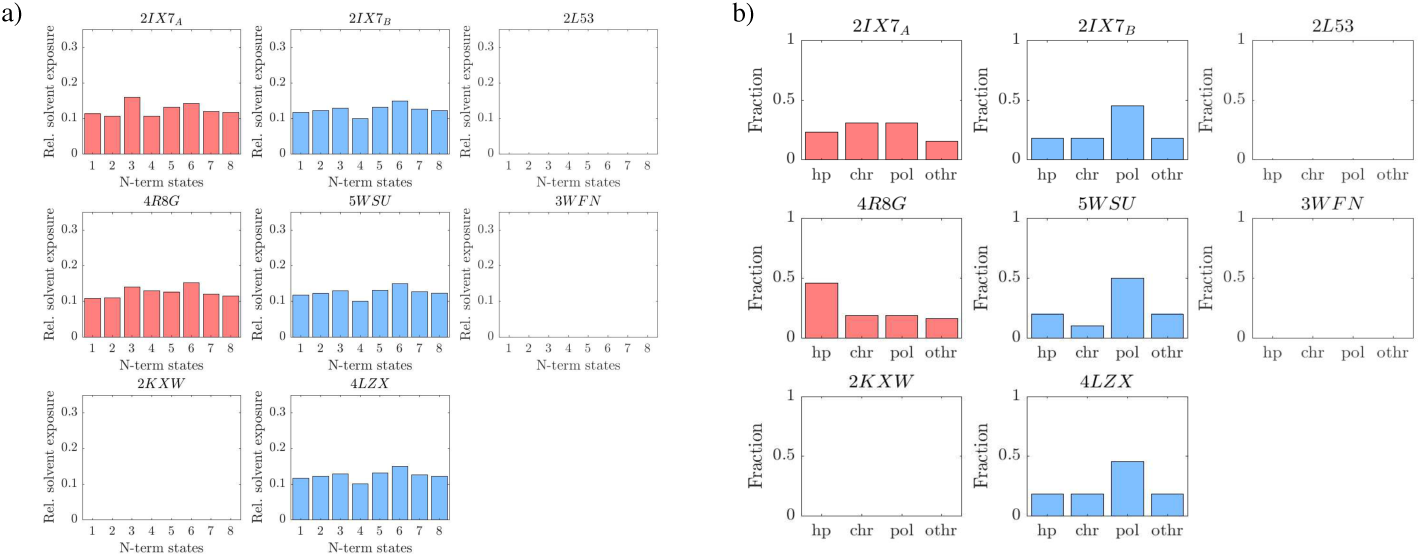
a) Apo N-CaM total relative solvent exposure and b) the residue types involved in apo N-CaM target protein contacts for each CaM-complex structure.

**S29 Fig.**
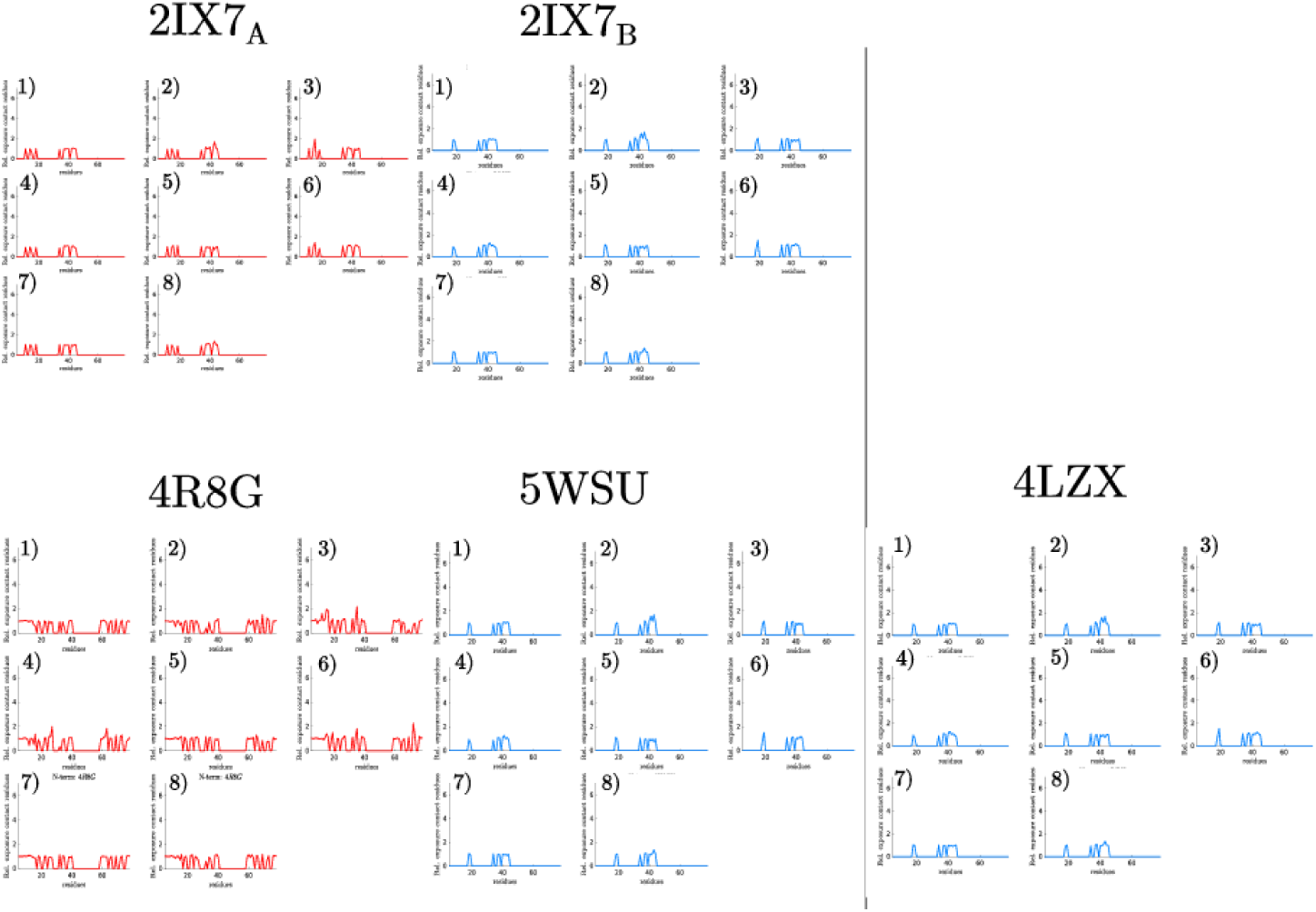
The relative solvent exposure per CaM-complex contact residue for the different apo N-CaM states.

S1 Data Interhelical angles, secondary structure assignments, and data used for solvent exposure analysis of all the states in holo and apo C-CaM and N-CaM.

S1 Text Explanation of interhelical angles and comparison of sampled states and experimental structures. A discussion of derived rate constants is also provided.

## Supplementary results

The representative structures of each state and a few similar experimental structures are shown in SI Fig 1-4. The holo C-CaM state 3, S1 Fig, corresponds to the helical arrangement found in 3CLN. Compared to this, state 2 has slightly smaller EH and larger EG angle, which correspond to a slight upward tilt of helix G and helix H C-term towards the linker. This is a similar helical arrangement as that of 5HIT. In addition to this, we note that interhelical angles are degenerate as descriptors for conformational states and thus their free energies. For example, holo N-CaM state 4 and 5SY1 C share the same ABC angles, S2 Fig, but display different arrangements of helices. Thus, contacts are better suited for clustering the conformational space (see methods).

Among the holo N-CaM representative structures, S2 Fig, we found states that strikingly resembled experimental structures. State 3 has similar helical arrangements as many experimental structures including 1CLL and 3CLN. This arrangement is characterized by a smaller CD angle, which results in the helices approaching a parallel arrangement. State 6, instead features an AC angle closer to perpendicular. Helix C is here almost parallel to the linker.

Among the apo C-CaM states, S3 Fig, state 2 is distinguished from the rest through smaller EG and larger GH angles. Here, helices F and G are placed in front of E and H, forming a compact conformation. In contrast to this, state 5 has a larger EG and smaller GH angle, which is the result of helix G facing away from E, opening the hydrophobic cleft.

The representative apo N-CaM structures, S4 Fig, display the resemblance between state 7 and 1DMO, where helix C is perpendicular to D, pointing in opposite directions, towards the reader. States 4, 6 and 8 have a larger AB angle than the other states. On these representative structures, the A and B helices are almost parallel, compared to states 3 and 5, where the helices A and B deviate more from being parallel. The main differences between state 8 and 4/6 are the angles between helices AC and CD. The orientation of helix C makes states 4 and 6 similar to 5WSU and 3WFN, while state 8 is not.

Rate constants derived from the MD simulations, S11 Fig, suggest that some states interconvert more rapidly than others. The rates *to* holo C-CaM state 2 are for example larger than rates *from* it, indicating a lower relative free energy. The same argument also applies to apo N-CaM state 6. Similarly, the rates suggest that apo C-CaM state 5 and holo C-CaM state 5 have higher relative free energies, and that holo N-CaM 4, 6, holo C-CaM 3, 6, apo N-CaM 4 and apo C-CaM 1 are separated by high free energy barriers.

## Acknowledgments

The simulations were performed on resources provided by the Swedish National Infrastructure for Computing (SNIC) at PDC Centre for High Performance Computing (PDC-HPC).

